# ACSS2 Regulates HIF-2α Degradation through the E3-Ubiquitin Ligase MUL1 in Clear Cell Renal Cell Carcinoma

**DOI:** 10.1101/2022.04.21.489116

**Authors:** Zachary A. Bacigalupa, Whitney A. Brown, Evan S. Krystofiak, Melissa M. Wolf, Rachel A. Hongo, Madelyn Landis, Edith K. Amason, Kathryn E. Beckermann, Jeffrey C. Rathmell, W. Kimryn Rathmell

## Abstract

Clear cell renal cell carcinoma (ccRCC) is an aggressive kidney cancer driven by *VHL* loss and aberrant HIF-2α signaling. Acetate metabolism may contribute to this axis by ACSS2-dependent acetylation of HIF-2α and may provide opportunities to intervention. Here we tested the effects of pharmacological and genetic manipulation of ACSS2 on HIF-2α, ccRCC cells, and tumors. ACSS2 inhibition led to HIF-2α degradation and suppressed ccRCC growth *in vitro*, *in vivo*, and in primary cell cultures of ccRCC patient tumors. This treatment resulted in reduced glucose and cholesterol metabolism, mitochondrial biogenesis and altered cristae deformation, that are consistent with loss of HIF-2α. Mechanistically, HIF-2α protein levels are regulated through proteolytic degradation and we found, in parallel to *VHL*, HIF-2α stability was dependent on ACSS2 activity to prevent direct interaction with the E3 ligase MUL1. These findings highlight ACSS2 as a critical upstream regulator of HIF-2α that may be exploited to overcome resistance to HIF-2α inhibitor therapies.

**STATEMENT OF SIGNIFICANCE:** We have unveiled ACSS2 as a critical upstream regulator of HIF-2α in ccRCC. Targeting ACSS2 potently promotes HIF-2α degradation via MUL1 to effectively deplete mitochondrial activity and block ccRCC primary tumor models and growth models resistant to HIF-2α inhibitor therapy.

## INTRODUCTION

Renal cell carcinoma (RCC) was diagnosed in 76,000 individuals in 2020 in the United States [1]. Clear cell renal cell carcinoma (ccRCC) is the most abundant and aggressive subtype of RCC, accounting for 75% of all RCC cases [2]. ccRCC has been intimately linked to the hereditary condition von Hippel-Lindau (VHL) disease, where patients harbor an inactivating mutation to the *VHL* tumor suppressor gene. In addition to familial associations, 90-95% of ccRCC cases are sporadically derived and the vast majority, upwards of 90%, present with a somatic mutation or suppressive hypermethylation of the promoter to the *VHL* gene [3–8]. The *VHL* gene encodes the E3 ubiquitin ligase pVHL, which functions to regulate the hypoxia inducible factor (HIF) family of transcription factors. Under normal oxygen conditions, prolyl hydroxylases (PHDs) add hydroxyl groups to proline residues on HIF-α proteins, which are then recognized by pVHL and targeted for proteasomal degradation [9]. Hypoxia, or the absence of pVHL, allows HIF-α proteins to accumulate, enter the nucleus and dimerize with HIF-1β/ARNT, and direct a major transcriptional response [10]. In ccRCC, HIF-2α is the major driver of transformation in part due to the loss of the 14q chromosomal arm where the HIF1A locus resides [11, 12]. Indeed, HIF-2α has been shown to directly regulate pathways such as lipid and cholesterol metabolism that fuel the clear cell phenotype [13–17]. Clinically, this has led to the development of small molecule inhibitors targeting HIF-2α interaction with the essential binding partner HIF-1β and HIF-2α transcriptional activity [18–21]. Belzutifan (MK-6482, previously PT-2399) is the first of such molecules to have received FDA approval for the treatment of adult patients with VHL disease who need therapy for ccRCC and other hemangioblastomas [22].

pVHL-mediated degradation of HIF-α proteins remains the most well understood mechanism of HIF-α disposal. However, VHL-independent degradation pathways have been identified, one involving JNK1-mediated stability of the chaperone proteins Hsp90/Hsp70, and another describes an endoplasmic reticulum (ER) stress-directed GSK3β-FBXW1A degradation axis [23]. More recently, the mitochondrial-tethered E3 ubiquitin ligase, MUL1, was found to indirectly promote HIF-1α degradation by targeting the HIF-1α degradation complex UBXN7 for proteasomal destruction [24]. Though MUL1 is best described to target proteins for degradation via SUMOylation, and K48 or K63 polyubiquitination, it can also aid mitophagy and autophagic degradation of mitochondrial associated proteins [25]. Localized to the outer mitochondrial membrane, MUL1 activity is integrated into mitochondrial quality control processes by modifying proteins such as DRP1 and Mitofusin 1, regulating the function and morphology of mitochondria [26, 27]. A role for MUL1 to regulate HIF-2α remains uncertain.

Targeting HIF-2α transcriptional activity has proven to be an effective treatment in VHL syndromic tumors, however, development of resistance makes parallel therapeutic strategies essential [28]. Resistance to belzutifan may be caused by HIF-2α mutations or the high levels of HIF-2α that can accumulate in the absence of pVHL and exceed local drug concentration. HIF-2α acetylation may provide an alternate approach to overcome resistance. Several studies have described the post-translational addition of an acetyl group to lysines on HIF-2α that supported protein stability and enhanced downstream signaling [29–32]. In these studies, the acetylation of HIF-2α, particularly during stress conditions, required the activity of acetyl-CoA synthetase 2 (ACSS2) [29–31]. ACSS2 is a metabolic enzyme present in both the cytoplasm and nucleus, which converts acetate into acetyl-CoA in an ATP-dependent reaction. In pre-clinical cancer models, ACSS2 has proven to be a promising target and has led to the rapid development of novel small molecule inhibitors [33–38]. ACSS2 has been found to facilitate epigenetic changes enhancing lysosomal biogenesis and autophagic activity in other biological models [39], which coupled with its reported interaction with physiologically expressed HIF-2α make ACSS2 a promising target in the context of ccRCC.

Here we investigate the relationship between ACSS2 activity and constitutive HIF-2α protein stability in the context of cancer models driven by *VHL* loss. We found that inhibition of ACSS2 resulted in transcriptional downregulation of genes involved in HIF-2α signaling, as well as cholesterol biosynthesis and glucose uptake. In addition, we found that ACSS2 inhibition induced HIF-2α proteasomal degradation even in the absence of pVHL. Mechanistically, ACSS2 inhibition or silencing stimulated the expression of MUL1, which formed a direct complex with HIF-2α to provide a potential VHL-independent and HIF2-α-specific mechanism to disrupt HIF-dependent transcription. Electron microscopy revealed ACSS2 inhibition, as well as HIF-2α silencing, caused massive mitochondrial deformation likely to result in catastrophic loss of function. Importantly, pharmacological inhibition of ACSS2 effectively blocked cancer cell growth in clinical samples and the HIF-2α inhibitor resistant 769-P cell line. Together, these findings demonstrate a critical role for ACSS2 to support ccRCC growth by interfering with direct regulation of HIF-2α stability by MUL1 to maintain the activity of key metabolic pathways. ACSS2 is, therefore, a novel therapeutic target for HIF-2α driven ccRCC.

## RESULTS

### ACSS2 is essential for ccRCC growth and proliferation

ACSS2 has remained largely unexplored as a potential therapeutic target in ccRCC. To examine the role of ACSS2 in ccRCC, we first performed a dose response, time course study with a pharmacological inhibitor of ACSS2 in HKC (normal kidney epithelia), 786-O (*VHL^-/-^* ccRCC), and A498 (*VHL^-/-^* ccRCC) cell lines. HKC cells were affected only at the highest dose, but ACSS2 inhibition led to a significant reduction in cell growth for the ccRCC cell lines (Figure 1A and 1B, Supplemental Figure 1A). We next analyzed the effect of ACSS2 inhibition on proliferation by measuring BrdU incorporation. Similarly, while HKC cells were only significantly affected after 48 hours of 5 µM and 10 µM doses, proliferation was significantly reduced in ccRCC cell lines at 24-hours with all doses of the inhibitor, (Figure 1C and 1D, Supplemental Figure 1B). Tumor sphere formation assays also revealed a striking difference in the ability of 786-O cells to form spheres when treated with ACSS2i (Supplemental Figure 1C). To assess the impact of ACSS2 on ccRCC growth more directly, we transduced the 786-O cell line to stably express a doxycycline-inducible shRNA construct targeting a control sequence or ACSS2. After 48 hours, we assessed cell growth via crystal violet staining and observed significant reduction in cell growth resulting from ACSS2 depletion (Figure 1E). Like the inhibitor, after 24 hours, 786-O cells expressing shACSS2 displayed significantly reduced BrdU incorporation (Figure 1F).

**Figure 1.**
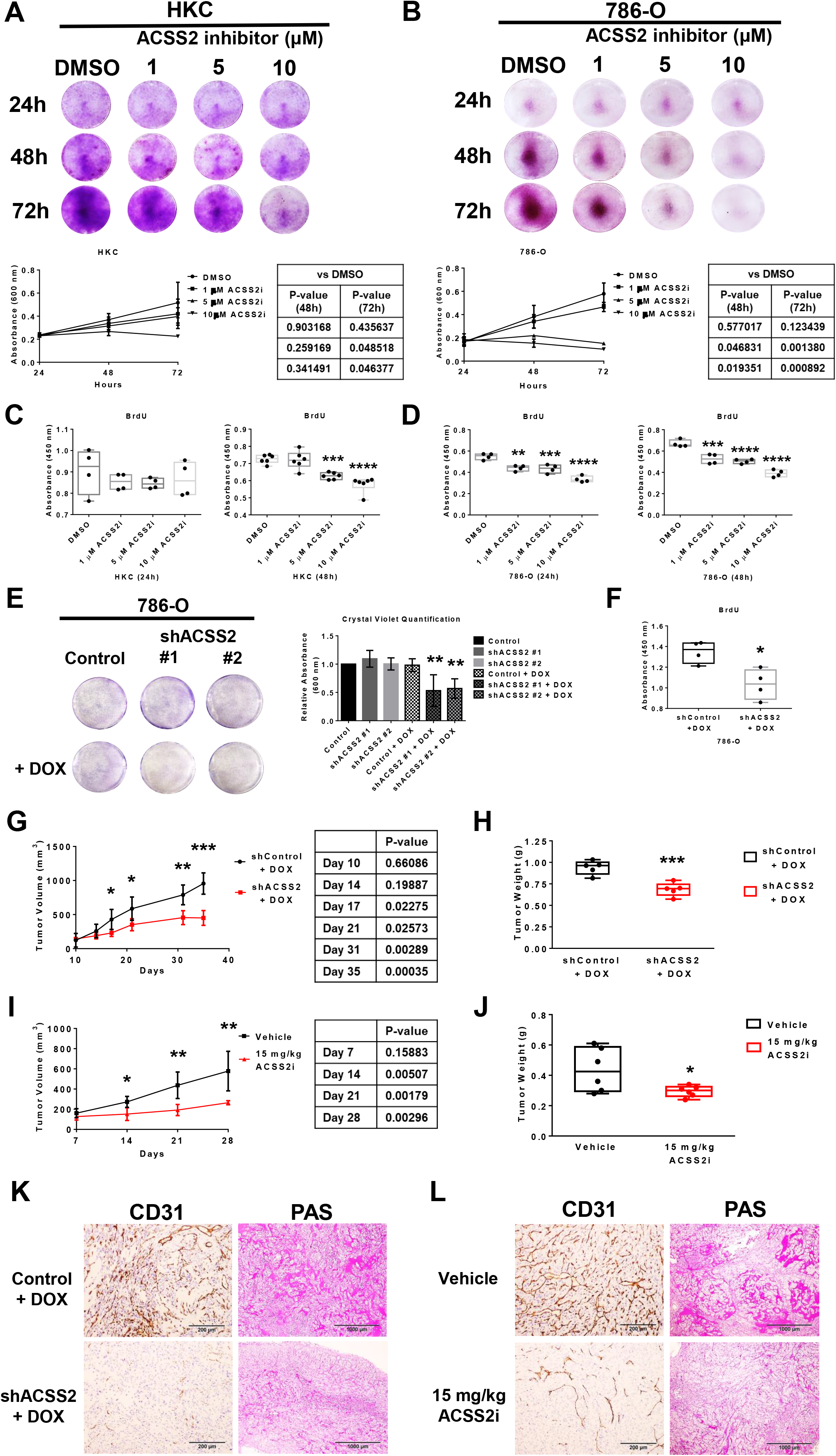
ACSS2 is essential for ccRCC growth and proliferation. **A.** Representative images of crystal violet staining from a dose-response time-course treatment with DMSO or ACSS2 inhibitor in HKC cells. Quantification of three-independent replicates are provided in the graph below. Statistical significance was determined using Bonferroni’s multiple comparisons test. **B.** Representative images of crystal violet staining from a dose-response time-course treatment with DMSO or ACSS2 inhibitor in 786-O cells. Quantification of three-independent replicates are provided in the graph below. Statistical significance was determined using Bonferroni’s multiple comparisons test. **C.** Box and whisker plots showing the absorbance values detected at OD450 nm of BrdU ELISA assays performed on HKC cells treated for 24 hours (left) or 48 hours (right) with DMSO, 1 µM, 5 µM, or 10 µM ACSS2 inhibitor (n=4). Statistical significance was determined using Bonferroni’s multiple comparisons test (**, P < 0.01; ****, P < 0.0001; n.s., not significant). **D.** Box and whisker plots showing the absorbance values detected at OD450 nm of BrdU ELISA assays performed on 786-O cells treated for 24 hours (left) or 48 hours (right) with DMSO, 1 µM, 5 µM, or 10 µM ACSS2 inhibitor (n=4). Statistical significance was determined using Bonferroni’s multiple comparisons test (**, P < 0.01; ***, P < 0.001; ****, P < 0.0001). **E.** Representative images of 786-O cells transduced with pTRIPZ Control (left), pTRIPZ shACSS2 #1 (center), or pTRIPZ shACSS2 #2 (right) untreated (top) or treated with 2 µg/mL doxycycline for 48 hours (bottom) stained with crystal violet. Bar graph displaying the quantification of crystal violet staining from three-independent experiments. Statistical significance was determined using a one-way ANOVA (**, P < 0.005). **F.** Box and whisker plots showing the absorbance values detected at OD450 nm of BrdU ELISA assays performed on 786-O pTRIPZ Control or 786-O pTRIPZ shACSS2 cells treated with 2 µg/mL doxycycline for 24 hours (n=4). Two-tailed paired t-test used to assess statistical significance (*, P<0.05). **G.** Line plot showing tumor volume over time in mice inoculated with 786-O pTRIPZ Control (black, n=5) or 786-O pTRIPZ shACSS2 (red, n=5) cells and fed a doxycycline rodent chow. Statistical significance was determined using multiple two-tailed t-tests. **H.** Box and whisker plot showing individual data points of tumor weight for pTRIPZ Control and pTRIPZ shACSS2 tumors. Statistical significance was determined using an unpaired, two-tailed t-test (***, P < 0.001). **I.** Line plot showing tumor volume over time in mice inoculated with 786-O cells treated with vehicle (black, n=6) or 15 mg/kg ACSS2 inhibitor (red, n=6). Statistical significance was determined using the Holm-Sidak method of multiple t-tests. **J.** Box and whisker plot showing individual data points of tumor weight for mice treated with vehicle (black, n=6) or 15 mg/kg ACSS2 inhibitor (red, n=6). Statistical significance was determined using an unpaired, two-tailed t-test (*, P < 0.05). **K.** Representative images taken at 10X magnification of sections from Control (top) and shACSS2 (bottom) tumors stained with CD31 (left) and Periodic Acid Schiff (PAS, right). **L.** Representative images taken at 10X magnification of sections from Vehicle (top) and 15 mg/kg ACSS2 inhibitor-treated (bottom) tumors stained with CD31 (left) and Periodic Acid Schiff (PAS, right).

With our data confirming ACSS2 is required for ccRCC growth in vitro and that ccRCC cell lines are more sensitive to ACSS2 inhibition, we next investigated the ability of ACSS2 inhibition to block tumor growth in vivo. Doxycycline induction of shACSS2 in mice bearing subcutaneous 786-O tumors resulted in a significant reduction in tumor growth (Figure 1G) and final tumor weight (Figure 1H). Similarly, mice bearing subcutaneous 786-O tumors receiving daily treatment with 15 mg/kg ACSS2i exhibited a remarkable decrease in tumor growth and final tumor weight (Figure 1I and 1J). In neither model did targeting ACSS2 result in any observable toxicity such as weight loss (Supplemental Figure 1D and 1E). Immunohistochemistry staining for CD31 and Periodic Acid Schiff revealed that targeting ACSS2 also reduced vascularization and glycogen deposition (Figure 1K-L and Supplemental Figure 1F-G), which are commonly associated clinical features of ccRCC linked to HIF signaling. Together, these data identify ACSS2 as a key contributor to tumor growth and a druggable vulnerability in ccRCC.

### HIF-2α gene and protein expression is regulated by ACSS2 activity

In ccRCC, aberrant HIF-2α signaling configures widespread signaling and metabolic changes. We hypothesized that these effects would be tied to the VHL-HIF axis. To test the role of ACSS2 to regulate essential metabolic signaling pathways in ccRCC, we performed transcriptomic analysis using the NanoString Metabolic Pathways probe set on 786-O cells transduced with shRNA targeting a control sequence or shACSS2 (Figure 2A). Intriguingly, differential expression analysis of this dataset identified a HIF-2α RCC Signaling gene set consisting of *EGLN3*, *EPAS1*, *PDGFB*, *SLC2A1*, and *VEGFA* as the most significantly downregulated pathway in response to ACSS2 deficiency (Figure 2B), suggesting that HIF-2α transcriptional activation was disrupted. To confirm these findings, we performed qRT-PCR on 786-O cells treated with 5 µM of ACSS2i for 24 hours (Supplemental Figure 2A) or transduced with shHIF2A (Supplemental Figure 2B) and observed a significant reduction in *Epas1* gene expression.

**Figure 2.**
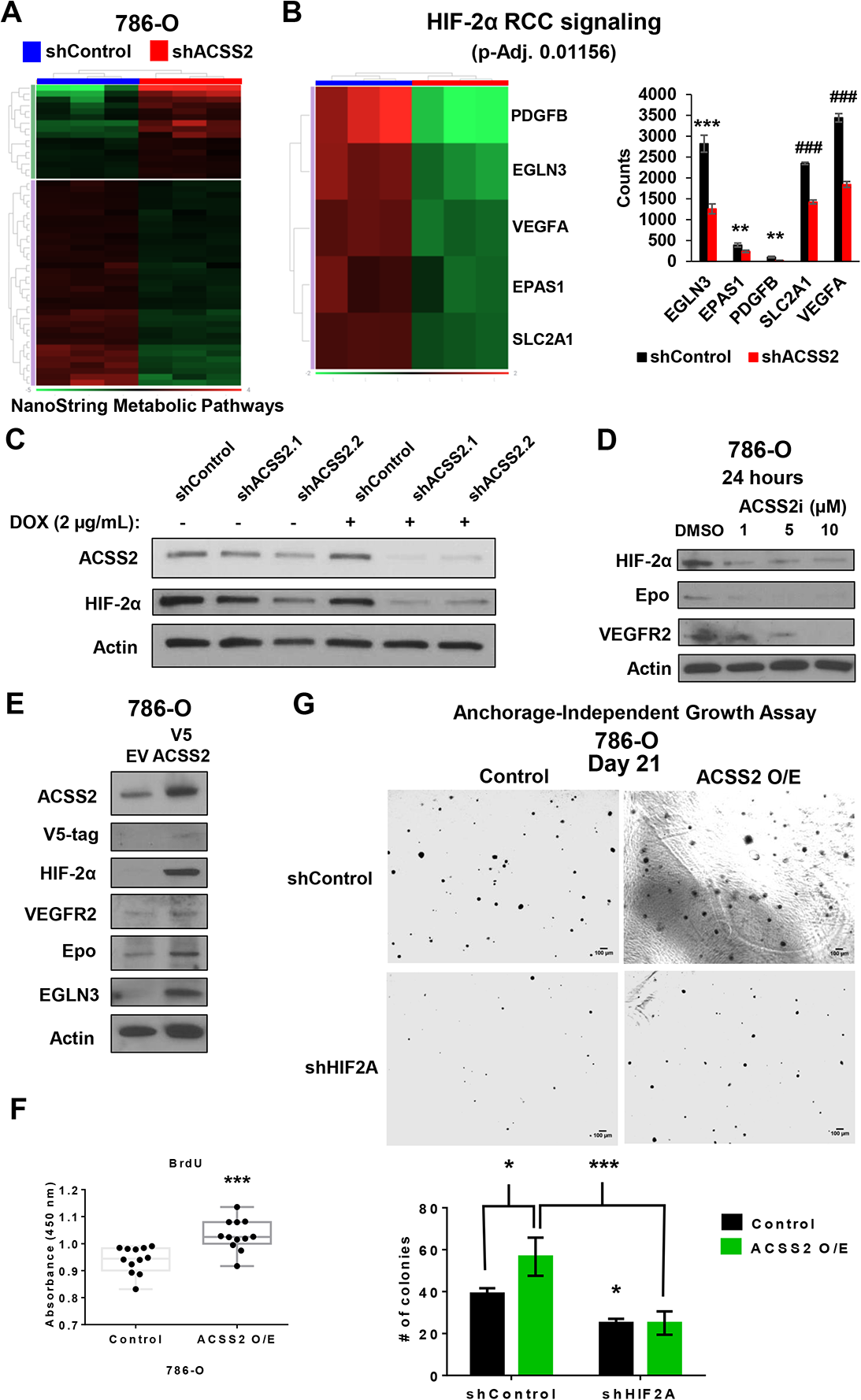
HIF-2α gene and protein expression is regulated by ACSS2 activity. **A.** Heatmap showing differential gene expression expressed as fold change (downregulated by –1.5 ≥ and upregulated by 1.5 ≤, p-Adj ≤ 0.05) between Control (blue) and shACSS2 (red) groups. Gene expression clustering on the Y-axis depicts genes that are upregulated (purple) and downregulated (green) in the Control group. **B.** Heatmap showing differential gene expression following pathway enrichment (Wikipathways) as fold change (downregulated by –1.5 ≥ and upregulated by 1.5 ≤, p-Adj ≤ 0.05) between Control (blue) and shACSS2 (red) groups. Gene expression clustering on the Y-axis depicts genes that are upregulated (purple) and downregulated (green) in the Control group. Bar graph (right) showing transcript counts from HIF-2α RCC Signaling pathway comparing Control (black) and shACSS2 (red) groups. Statistical significance determined using unpaired t-tests without correction for multiple comparisons (**, P < 0.01; ***, P < 0.001; ###, P < 0.0001). **C.** Western blot analysis of HIF-2α and ACSS2 expression following 24-hour doxycycline induction of 786-O pTRIPZ Control and pTRIPZ shACSS2 cells. **D.** Western blot analysis assessing expression of HIF-2α, Epo, and VEGFR2 in 786-O cells treated with DMSO, 1 µM, 5 µM, or 10 µM ACSS2 inhibitor for 24 hours. **E.** Western blot analysis showing expression of ACSS2, V5-tag, HIF-2α, VEGFR2, Epo, and EGLN3 in 786-O cells transduced pLX304 empty vector or pLX304 V5-ACSS2 overexpression vector. **F.** Box and whisker plot showing the absorbance values detected at OD450 nm of BrdU ELISA assays performed on 786-O cells transduced pLX304 empty vector or pLX304 V5-ACSS2 overexpression vector. Statistical significance was determined using an unpaired, two-tailed t-test (***, P < 0.001). **G.** Representative images taken at day 21 of an anchorage-independent growth assay performed using 786-O control or V5-ACSS2 cells transduced express shControl or shHIF2A. Bar graph showing the number of colonies formed for each condition from three-independent experiments. Statistical significance determined using two-way ANOVA and Tukey’s multiple comparisons test (*, P-value < 0.05; ***, P-value < 0005).

Because HIF-2α is largely regulated at the protein level, we hypothesized ACSS2 may impact HIF-2α protein. Consistent with this model, we observed a robust decrease in HIF-2α protein in 786-0 cells transduced with shACSS2 (Figure 2C). HIF-2α protein expression was also suppressed by pharmacologic ACSS2 inhibition to result in reduced expression of known HIF-2α targets (Figure 2D). Notably, HIF-2α protein expression was not impacted by ACSS2 inhibition in the HKC cell line (Supplemental 2C). To further test the involvement of ACSS2 activity for the observed changes, we stably overexpressed ACSS2 in 786-O cells and assessed protein expression, where we observed an increase in the expression of HIF-2α and its downstream signaling network (Figure 2E). Additionally, a BrdU proliferation assay confirmed that overexpression of ACSS2 in 786-O cells also resulted in enhanced proliferation (Figure 2F). Next, we tested whether overexpressing ACSS2 was sufficient to enhance tumorigenic potential and determine if any observed effect was dependent on HIF-2α utilizing an anchorage-independent growth assay. 786-O cells were transduced to express V5-ACSS2 or the empty vector followed by a second transduction to target HIF2A with shRNA (Supplemental Figure 2D). After 21 days of growth, it was determined that ACSS2 overexpression increased the size and number of colonies, but this increase was fully mitigated when *HIF2A* was targeted with shRNA (Figure 2G).

### ACSS2 activity mediates HIF-2α stability

Previous studies have identified acetylated lysine residues on HIF-2α, which confer protein stability and yield optimal signaling, that are acquired from acetyl-CoA produced from ACSS2 [29–32]. To better understand the mechanism by which ACSS2 may regulate HIF-2α protein expression, we performed a 1-hour incubation with the proteasomal inhibitor MG132 followed by the addition of DMSO or 5 µM ACSS2i for 24 hours. Interestingly, 786-O cells receiving the MG132 pre-treatment prior to ACSS2i displayed a significant rescue in HIF-2α expression (Figure 3A and Supplemental 3A) suggesting ACSS2 inhibition enhances HIF-2α proteasomal degradation, even in *VHL* deficient 786-O cells.

**Figure 3.**
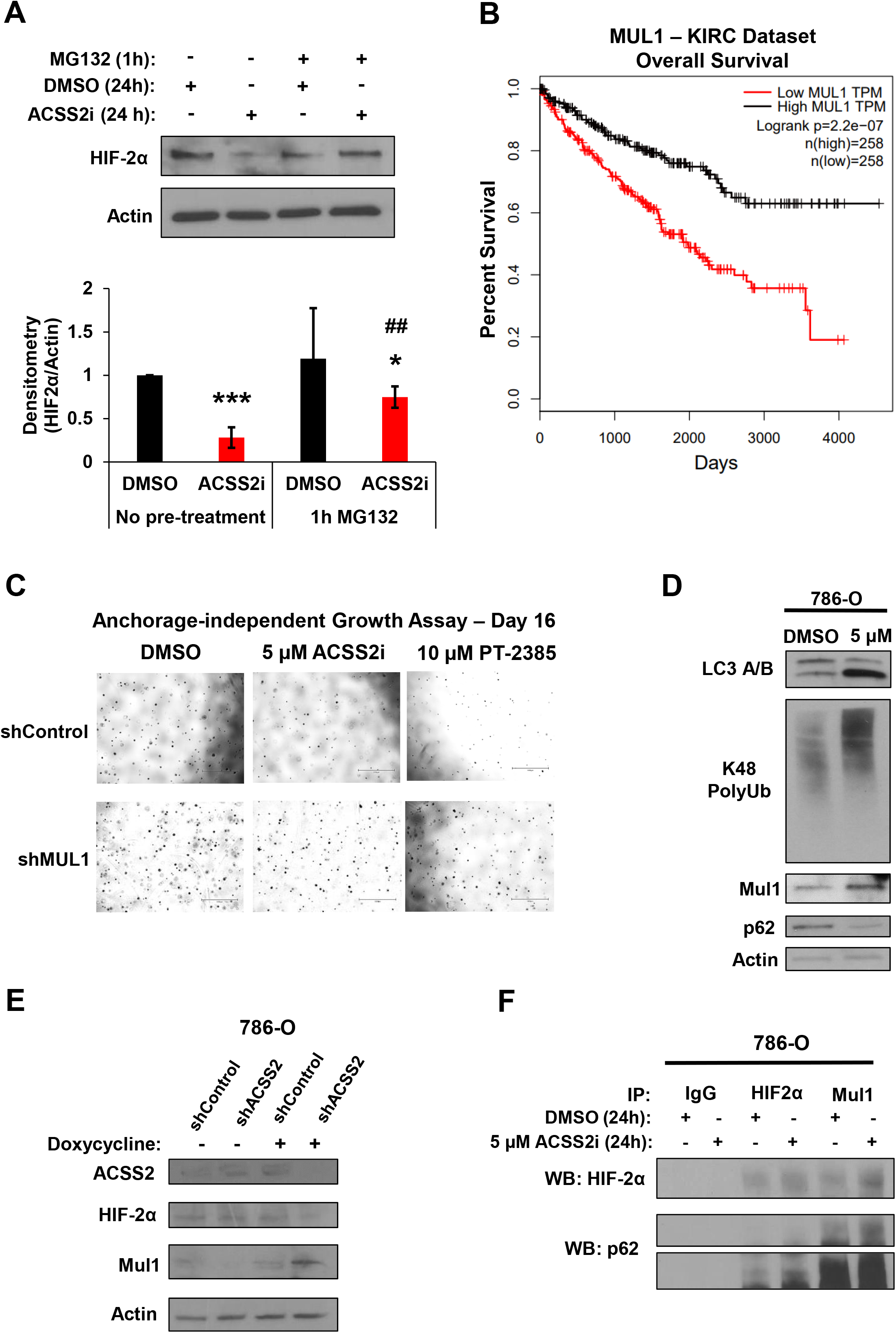
ACSS2 activity mediates HIF-2α stability. **A.** Western blot depicting HIF-2α expression in 786-O cells in the absence or presence of a 1-hour incubation with 10 µM MG132 prior to being treated with DMSO or 5 µM ACSS2 inhibitor for 24 hours. Bar chart (right) showing densitometry analysis performed using ImageJ on western blot images from three-independent experiments. Statistical significance determined using multiple unpaired, two-tailed t-tests (*, P < 0.05; ***, P < 0.0005; ##, P < 0.01; * = t-test performed vs No pre-treatment DMSO; # = t-test performed vs No pre-treatment ACSS2i). **B.** Kaplan-Meier plot generated from the KIRC TCGA dataset assessing patient outcome as it relates to MUL1 expression (MUL1^HIGH^ = black, MUL1^LOW^ = red). Statistical significance determined automatically via GEPIA database. **C.** Representative images at day 16 of anchorage-independent growth assays performed on 786-O cells transduced to express shControl or shMUL1 and treated with either DMSO, 5 µM ACSS2 inhibitor, or 10 µM PT-2385. **D.** Western blot images showing expression of the mitophagy marker LC3 A/B, K48 polyubiquitination, MUL1, and p62 in 786-O cells treated with DMSO or 5 µM ACSS2 inhibitor for 48 hours. **E.** Western blot analysis showing expression changes to ACSS2, HIF-2α, and MUL1 in response to induction of shACSS2. **F.** Western blot images assessing protein interactions from immunoprecipitations of HIF-2α or MUL1 performed on 786-O cells treated with DMSO or 5 µM ACSS2 inhibitor for 24 hours.

Recently, several studies have linked the E3 ubiquitin ligase, MUL1, to the indirect regulation of HIF-1α degradation by the E3-ligase co-factor UBXN7 [24]. Consistent with a potential role for MUL1, analysis of TCGA and CPTAC samples via the UALCAN database show that MUL1 transcript abundance and protein expression are significantly reduced when compared to normal adjacent tissue (Supplemental Figure 3B and 3C). Survival analysis performed on the KIRC dataset assessing the effect of MUL1 expression on patient outcome identified a correlation between low MUL1 expression and reduced survival (Figure 3B). Intriguingly, depleting MUL1 via shRNA provided 786-O cells with resistance to treatment with ACSS2i or PT-2385, enabling successful anchorage-independent growth (Figure 3C and Supplemental Figure 3D). Considering these findings, we next analyzed the effect of targeting ACSS2 on MUL1 expression. Pharmacologically or genetically targeting ACSS2 in 786-O cells increased MUL1 expression (Figure 3D and 3E). In the HKC cell line, we also observed an increase in MUL1 expression in response to ACSS2 inhibition despite having no impact on HIF-2α expression, which was possibly due to continued expression of pVHL (Supplemental Figure 3E). To test whether MUL1 could interact with and direct the degradation of HIF-2α, we immunoprecipitated MUL1 from 786-O cells treated with DMSO or 5 µM ACSS2i for 24 hours and observed HIF-2α and MUL1 to interact in a complex (Figure 3F). These results show that ACSS2 is a critical and direct mediator of MUL1-directed HIF-2α protein degradation. Moreover, these findings highlight ACSS2 and MUL1 as means to target HIF-2α in ccRCC.

### ACSS2 activity supports glucose and cholesterol metabolism

The link between hypoxia signaling and metabolism has been well-characterized and aberrant HIF-2α signaling has been shown to promote glucose and cholesterol metabolism in ccRCC. To address the impact of ACSS2 on metabolic pathways, we performed a transcriptomic analysis. Differential gene expression analysis of 786-O cells treated with DMSO or 5 µM ACSS2i for 24 hours identified cholesterol metabolism as the most significantly altered pathway (Figure 4A). Notable changes within the cholesterol metabolism pathway corresponding to ACSS2 inhibition included the decrease in HMG-CoA synthase (HMGCS1), which synthesizes HMG-CoA from acetyl-CoA, HMG-CoA reductase (HMGCR), whose catalytic activity produces mevalonate in the rate-limiting step of cholesterol synthesis, and 7-dehydrocholesterol reductase (DHCR7) and 24-dehydrocholesterol reductase (DHCR24) which perform the final reaction in cholesterol biosynthesis (Supplemental Figure 4A). Functionally, we measured total cholesterol levels in 786-O cells treated with DMSO or 5 µM ACSS2i for 24 hours and found a significant reduction in response to ACSS2 inhibition (Figure 4B).

**Figure 4.**
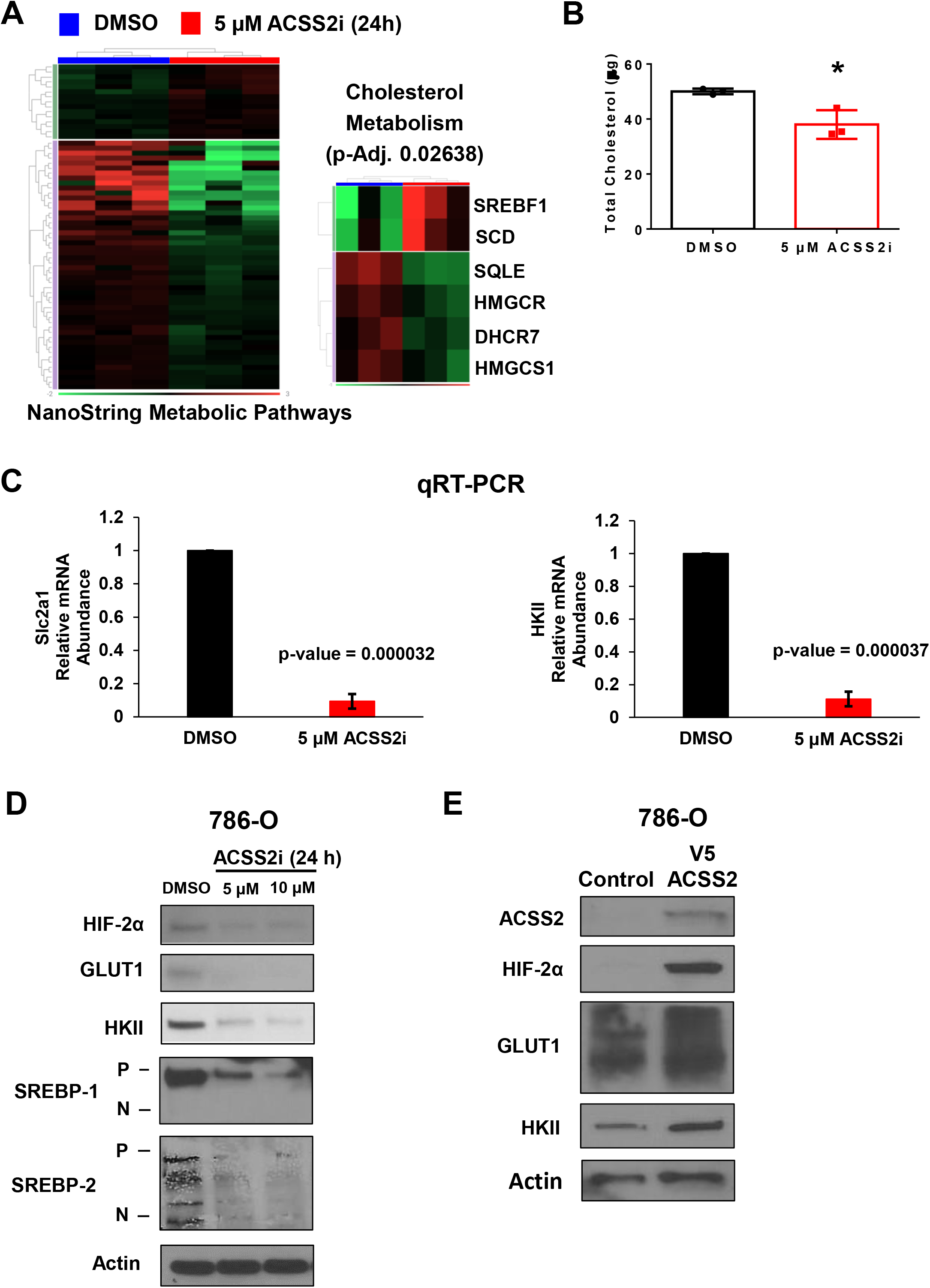
ACSS2 activity supports glucose and cholesterol metabolism. **A.** Heatmap (left) showing differential gene expression expressed as fold change (downregulated by –1.1 ≥ and upregulated by 1.1 ≤, p-Adj ≤ 0.05) between 24-hour treatment of 786-O cells with DMSO (blue) and 5 µM ACSS2 inhibitor (red) groups. Gene expression clustering on the Y-axis depicts genes that are upregulated (purple) and downregulated (green) in the DMSO group. Heatmap (right) showing differential gene expression following pathway enrichment (Wikipathways) as fold change (downregulated by –1.1 ≥ and upregulated by 1.1 ≤, p-Adj ≤ 0.05) between 24-hour treatment of DMSO (blue) and 5 µM ACSS2 inhibitor (red) groups. Gene expression clustering on the Y-axis depicts genes that are upregulated (purple) and downregulated (green) in the DMSO group. **B.** Bar graph showing total cholesterol levels (µg) in 786-O cells treated with DMSO (black; n=3) or 5 µM ACSS2i (red; n=3) for 24 hours. Statistical significance determined using an unpaired, two-tailed Student’s t-test. **C.** Bar graphs showing relative mRNA abundance of SLC2A1 (left) and HKII (right) in 786-O cells following 24-hour treatment with DMSO (black) or 5 µM ACSS2 inhibitor (red). Statistical significance determined using an unpaired, two-tailed Student’s t-test. **D.** Western blot analysis showing expression of HIF-2α, GLUT1, HKII, SREBP-1, and SREBP-2 in 786-O cells treated with DMSO, 5 µM, or 10 µM ACSS2 inhibitor for 24 hours. **E.** Western blot analysis of ACSS2, HIF-2α, GLUT1, and HKII protein expression in 786-O cells transduced to express pLX304 empty vector or pLX304 V5-ACSS2 overexpression vector.

Given the link between hypoxia signaling and glucose metabolism, we next investigated the effect of ACSS2 inhibition on *SLC2A1* (Glut1) and hexokinase II (*HKII*) gene expression. To this end, we performed RT-PCR on 786-O cells treated with DMSO or 5 µM ACSS2i for 24 hours and found the transcripts of *SLC2A1* and *HKII* to be significantly reduced in response to ACSS2 inhibition (Figure 4C). Next, we tested if the observed mRNA changes correspondingly altered protein expression. Following 24 hours of treatment with DMSO or ACSS2i, HIF-2α, as well as GLUT1, HKII, SREBP-1, and SREBP-2 which are more commonly associated with HIF-1α [40, 41], protein levels were all decreased in response to ACSS2 inhibition (Figure 4D). Conversely, overexpression of ACSS2 in 786-O cells enhanced expression of HIF-2α, GLUT1, and HKII (Figure 4E). A time course measurement on glucose present in the media of 786-O cells treated with DMSO or 5 µM ACSS2i revealed a significant increase in glucose present in the media after 48 hours of ACSS2i treatment, indicating an impaired ability of the cells to uptake and consume glucose (Supplemental 4B). Additionally, we quantified the acetate concentrations in the media of 786-O cells following a 24-hour dose response with ACSS2i and found that ACSS2 inhibition significantly enhanced the amount of acetate present in the media (Supplemental Figure 4C). Similarly, 786-O cells transduced with the doxycycline-inducible shRNA system saw an increase in acetate in the media when shACSS2 was induced, whereas a significant decrease in acetate was observed when ACSS2 was overexpressed (Supplemental Figure 4D). These results confirm that altering ACSS2 activity directly impacts acetate consumption. Furthermore, our results demonstrate that ACSS2 activity transcriptionally regulates the rate-limiting enzymes in cholesterol synthesis and glucose metabolism, which are likely to have widespread effects on the cell.

### Targeting ACSS2 elicits cancer cell-specific mitochondrial defects

Mitochondrial quality is intimately linked to their function and ability to facilitate metabolic processes [42, 43]. Considering the magnitude by which targeting ACSS2 has shown to inhibit glucose metabolism and cholesterol biosynthesis, we hypothesized that inhibiting ACSS2 would selectively disrupt mitochondrial homeostasis in cancer cells. To test the effects on mitochondrial morphology electron microscopy was performed on HKC and 786-O cells treated with DMSO for 72 hours or with 5 µM ACSS2i for 24, 48, and 72 hours (Figure 5A and 5B). Strikingly, the mitochondria in 786-O cells treated with ACSS2i began to transform as early as 24 hours from the typically observed elongated morphology as observed in the DMSO control, to a smaller and spherical morphology, a process not detected in the mitochondria of HKC cells (Figure 5A and 5B). Additionally, the cristae morphology degenerated from linear and continuous to fragmented and spherical (Figure 5B). These results were further corroborated when we looked at the expression of proteins involved in cristae structure such as Opa-1, Mitofusin 1, Mitofusin 2, and Drp1, after treatment with 5 µM ACSS2i for 48 hours result in a loss of expression specifically in 786-O cells (Supplemental 5A and 5B). We also observed a cancer cell-specific reduction in the expression of PGC1β, which has been shown to control mitochondrial biogenesis [44] (Figure 5A and 5B).

**Figure 5.**
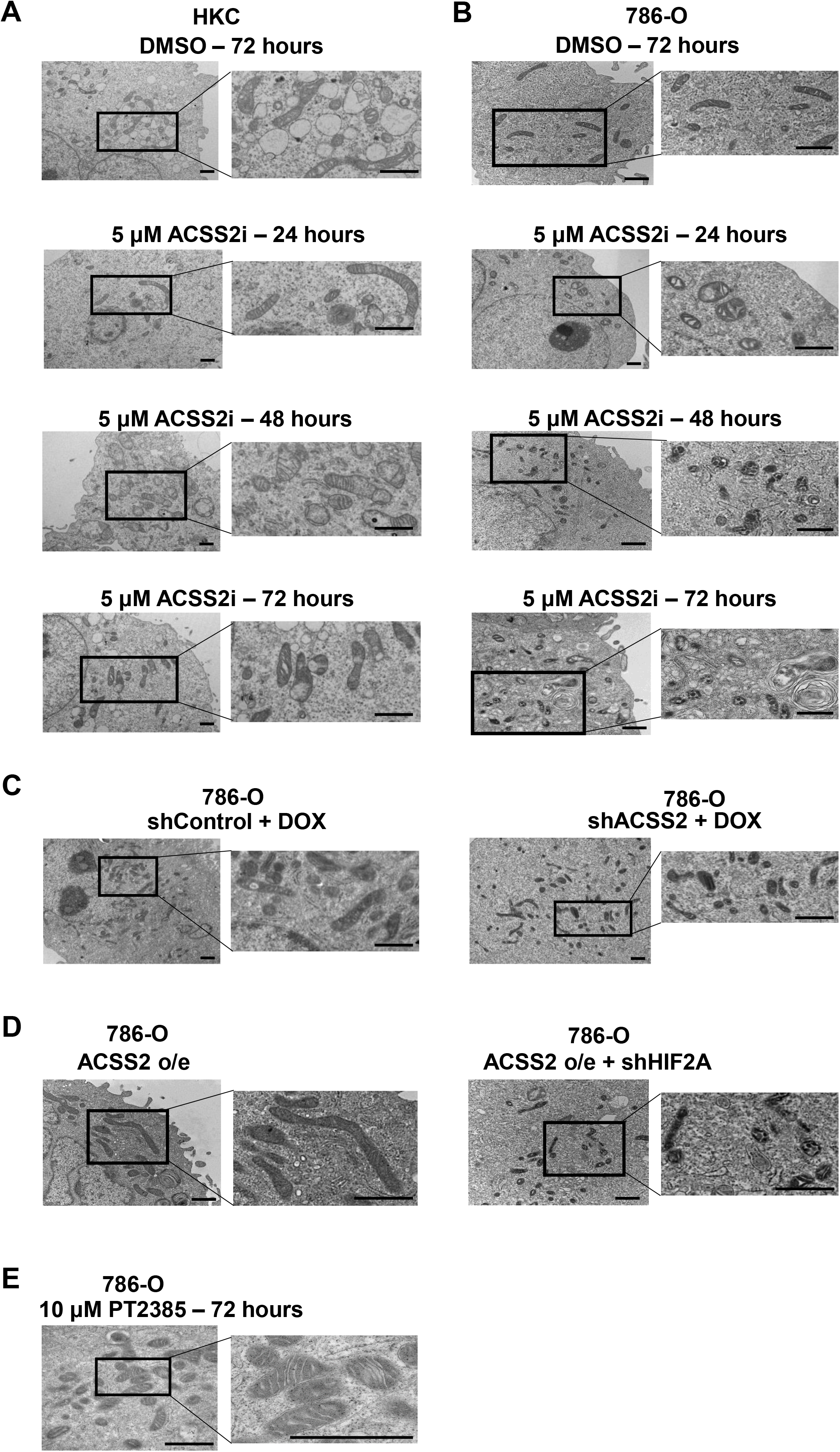
Targeting ACSS2 elicits cancer cell-specific mitochondria morphology defects observed by transmission electron microscopy. Representative images of HKC control cell mitochondria (**A**) and 786-O mitochondria (**B**) treated with DMSO for 72 hours or with 5 µM ACSS2 inhibitor for 24, 48, and 72 hours. **C.** 786-O pTRIPZ Control and shACSS2 cells induced with 2 µg/mL doxycycline for 48 hours. **D.** 786-O cells transduced to overexpress ACSS2 (left) followed by transduction with shHIF2A (right). **E.** 786-O cells treated with 10 µM PT-2385 for 72 hours. Black square insets indicate areas of higher magnification shown to the right of the images. Scale bars are 1 µm for all images.

The impact of ACSS2 inhibition on mitochondrial biogenesis was next tested. Applying a colorimetric assay that detects both the nuclear encoded mitochondrial gene SDH-A and the mitochondrial encoded COX-I, we discovered a significant, dose-dependent decrease in COX-I after 24 hours of ACSS2i treatment in the 786-O cells. This had no impact, however, on the HKC cells (Supplemental 5C and 5D). To determine if the observed mitochondrial phenotype was specific to ACSS2, we performed electron microscopy on 786-O cells where shRNA targeting ACSS2 had been induced with doxycycline for 48 hours and we observed shACSS2 induction had the same morphological changes that pharmacological inhibition produced (Figure 5C). Additionally, shACSS2 had a similar effect on mitochondrial biogenesis (Supplemental Figure 5E). However, the reduced expression of cristae structural proteins observed in response to ACSS2i was not as robust with shACSS2, although PGC1β induction was lost (Supplemental Figure 5F). Together, these data suggest that ACSS2 activity serves an integral role to maintain the quality of mitochondria in *VHL*-mutated cancer cells, possibly through a PGC1β-dependent mechanism.

We next questioned whether mitochondrial maintenance influenced by ACSS2 activity was dependent on HIF-2α. To address this question, electron microscopy was performed on 786-O cells overexpressing ACSS2 which had been transduced to express shControl or shHIF2A. Mitochondria from cells overexpressing ACSS2 had a characteristically normal appearance displaying an elongated morphology with defined cristae; however, when shHIF2A was introduced, we observed a return of the small, circular mitochondrial phenotype with deformed cristae (Figure 5D). Interestingly, treatment with the HIF-2α inhibitor PT-2385 did not result in a morphological defect in the mitochondria (Figure 5E), possibly indicating that PT-2385 has lower efficacy than targeting with shRNA. In sum, these results suggest a role for HIF-2α in maintaining healthy mitochondria and provide further insight into the scope of the ACSS2-HIF-2α axis.

### ACSS2 inhibition selectively impedes cancer cell growth and HIF-2α expression in ccRCC patient samples

The HIF-2α inhibitor belzutifan was shown to be highly effective in patients with von Hippel-Lindau disease and has recently gained FDA approval for the treatment of adult patients with the disease who need treatment for RCC. As our previous results demonstrated that targeting ACSS2 is an effective strategy to selectively target HIF-2α, we next moved our experiments into primary cell cultures isolated from ccRCC patients. A time-course, dose-response treatment was performed on matching normal adjacent tissue and cancer cells from three patients and cancer cells again observed to be significantly more sensitive than normal epithelial cells to ACSS2 inhibition (Figure 6A-B and Supplemental Figure 6A-D). Next, we explored the effect of ACSS2 inhibition on HIF-2α expression and found that 24 hours of treatment at all doses of ACSS2i in these clinical specimens was effective at reducing HIF-2α expression in a cancer cell-specific manner (Figure 6C and 6D). Additionally, HKII expression increased and both findings were supported by transcriptomic analysis to corroborate our previous findings that ACSS2 inhibition reduces the expression of genes involved in cholesterol biosynthesis (Figure 6C-D and Supplemental Figure 6E-F). Lastly, we explored whether ACSS2 inhibition could be an effective means to treat ccRCC that are resistant to HIF-2α inhibitor therapy. To this end, we performed a dose-response, time-course ACSS2i treatment in the previously described HIF-2α inhibitor resistant 769-P ccRCC cell line and found that ACSS2i, when provided as single agent, was able to block the growth of these cells (Figure 6E). Interestingly, a slight synergistic effect was observed when treating 769-P cells with the ACSS2 inhibitor and PT-2385, suggesting that providing ACSS2i in conjunction with HIF-2α inhibition could be effective at circumventing resistance to deliver a more effective suppression of HIF-2α driver signals (Figure 6E). Altogether, these results demonstrate that our in vitro and in vivo studies translate to ccRCC clinical samples, highlighting ACSS2 as a novel way to target HIF-2α and major metabolic pathways specifically in cancer cells.

**Figure 6.**
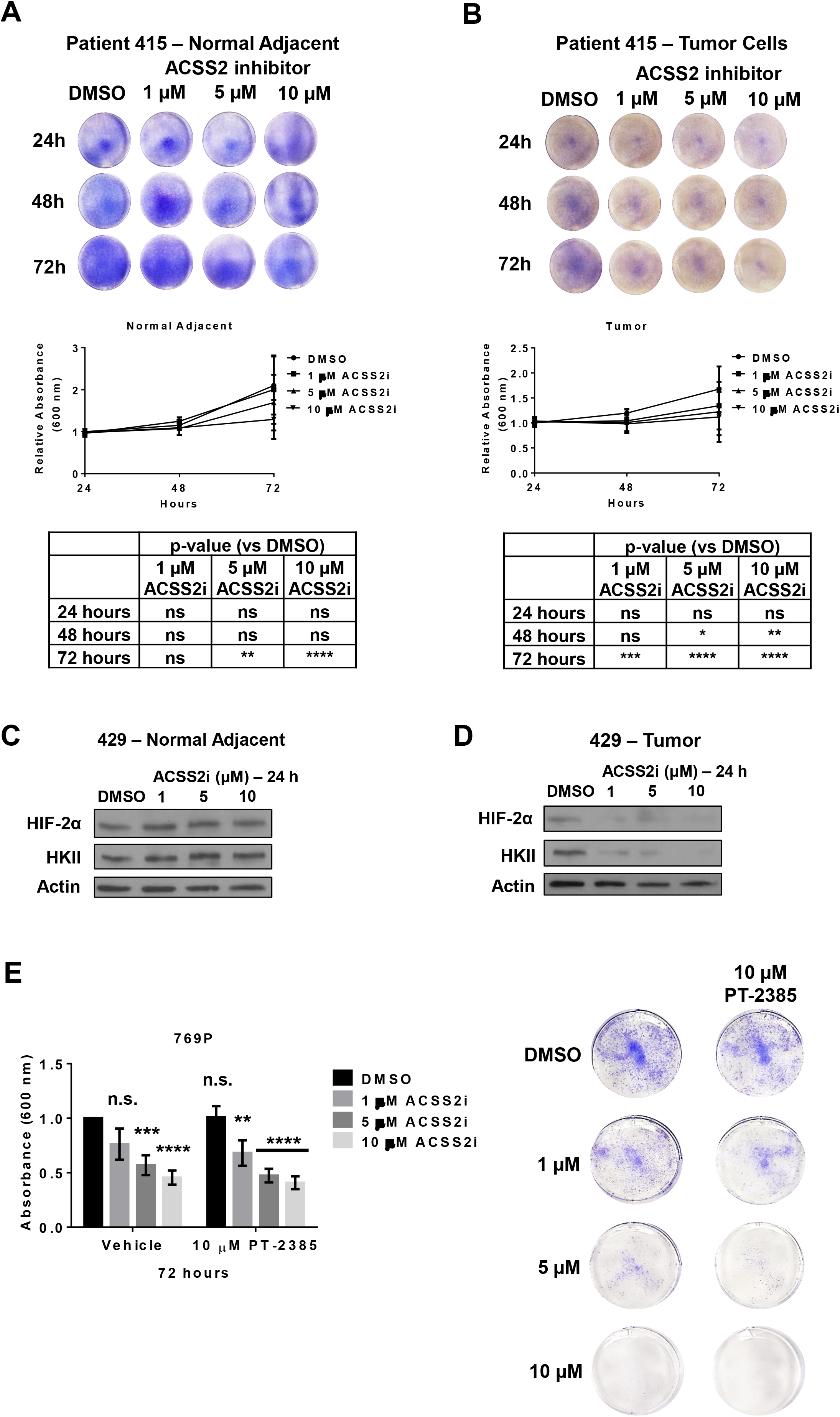
ACSS2 inhibition selectively impedes cancer cell growth and HIF-2α expression in ccRCC patient samples. **A.** Representative images of crystal violet staining from a dose-response time-course treatment with DMSO or ACSS2 inhibitor in epithelial cells isolated from ccRCC patient’s normal adjacent tissue. Quantification of three-independent replicates are provided in the graph below. Statistical significance was determined using Bonferroni’s multiple comparisons test (P values provided in table below). **B.** Representative images of crystal violet staining from a dose-response time-course treatment with DMSO or ACSS2 inhibitor in epithelial cells isolated from ccRCC patient tumor sections. Quantification of three-independent replicates are provided in the graph below. Statistical significance was determined using Bonferroni’s multiple comparisons test (P values provided in table below). **C.** Representative western blot images showing expression of HIF-2α and HKII in cells isolated from ccRCC patient normal adjacent tissue treated with DMSO, 1µM, 5 µM, or 10 µM ACSS2 inhibitor for 24 hours. **D.** Representative western blot images showing expression of HIF-2α and HKII in epithelial cells isolated from ccRCC patient tumor sections treated with DMSO, 1µM, 5 µM, or 10 µM ACSS2 inhibitor for 24 hours. **E.** Bar graph showing crystal violet quantification of 769-P cells (three-independent experiments) in the presence or absence of 10 µM PT-2385 treated with DMSO, 1µM, 5 µM, or 10 µM ACSS2 inhibitor for 24, 48, and 72 hours. Statistical significance determined using a two-way ANOVA and Tukey’s multiple comparisons test. Representative images of crystal violet staining from a 72-hour dose-response treatment with DMSO or ACSS2 inhibitor in 769-P cells.

## DISCUSSION

ACSS2 has recognized functions in the dynamic interplay of stress responses, including interaction with HIF-2α in the setting of adaptive responses to oxygen and nutrient limited states [29, 30, 35, 36, 45]. In clear cell renal cell carcinoma, the pseudohypoxic state is defined by high level, constitutive HIF-2α signaling that is unimpeded by normal stress signals. We sought to directly interrogate the interaction of ACSS2-mediated acetylation to maintain the stability of HIF-2α in the absence of *VHL* and provide a key driver of these tumors [46–49]. Recent work has advanced a HIF-2α specific transcriptional inhibitor for treatment of *VHL*-associated renal cell carcinomas although resistance can develop over time[18, 50]. Parallel strategies to block the HIF-2α node, such as targeting alternate mechanisms of HIF-2α protein stability will be necessary for adequate disease control.

By applying various strategies to alter ACSS2 activity, we showed that ccRCC cells have some level of dependence on ACSS2 to support HIF-2α expression, stability, and signaling, which makes them inherently more vulnerable to ACSS2 inhibition. Importantly, we also established that ACSS2 inhibition was effective to block the growth of cancer cells derived from ccRCC patients, as well as the HIF-2α inhibitor resistant 769-P cell line. Our data suggest that treatment with an ACSS2 inhibitor could have a potent effect at treating ccRCC patients who have a *VHL* mutation and could possibly have a synergistic effect when used in combination with HIF-2α inhibition. Novel small molecules targeting ACSS2 are in development [34] and being characterized in preclinical models of cancer. Indeed, while HIF-2α inhibition with belzutifan has been revolutionary for VHL syndrome, the effect is slow. Also, although each cancer in these scenarios displays HIF-2a activation and transcriptional dependencies, simple inhibition of dimerization does not promote cell death promptly. We show here that acetylation via ACSS2 is important for HIF-2α stabilization, that inhibiting ACSS2 reduces transcription of EPAS1 as well as induces protein degradation via a secondary mechanism. This more potent blockade effectively shuts down hypoxia transcription and selectively impairs mitochondrial function in a way that proves to be lethal to cells that are dependent on HIF-2α signaling.

*Vhl* is mutated in over 70% of ccRCC and aberrant HIF-2α signaling persists due to a lack of an effective degradation mechanism. The identification and characterization of alternative HIF-2α degradation pathways could manifest into clinically relevant findings. Recently, the mitochondria-associated E3 ubiquitin ligase MUL1 has been shown to indirectly regulate the degradation of HIF-1α through its action of UBXN7, and was also found to promote autophagy and be suppressed in ccRCC cancer cells [24, 51]. Given the high level of homology between HIF-1α and HIF-2α and the link made to MUL1 and ccRCC, we rationalized interrogating a link between ACSS2 and MUL1 as a mechanism to alter HIF-2α stability. Identifying that MUL1 directly interacts with HIF-2α and that ACSS2 inhibition can enhance MUL1 expression and its interaction with HIF-2α, implicates MUL1 as a potential mechanistic link as to how ACSS2 activity regulates HIF-2α in a tumor lacking *Vhl* (Supplemental Figure 6G). Beyond direct polyubiquitin-mediated degradation, it has also been demonstrated that MUL1 can contribute to the degradation of targets such as Mitofusin 1 via mitophagy [25, 27]. Interestingly, it was recently shown that metabolic stress induces the nuclear localization of ACSS2, where it promotes acetylation and interacts with TFEB to facilitate the transcription of genes involved in lysosomal biogenesis and autophagy [39]. Taking these findings into consideration with our own, it would be intriguing to investigate the involvement of mitophagy in the ACSS2-dependent degradation of HIF-2α.

Acetate is a major precursor to produce acetyl-CoA, particularly under times of metabolic stress. While citrate-derived acetyl-CoA has been shown to be more heavily utilized in lipid synthesis [52, 53], acetyl-CoA produced from ACSS2 is thought to have more of a role in post-translational and epigenetic regulation [29, 31, 38]. Though the findings we describe here remain focused on the relationship between ACSS2 and HIF-2α, the role of ACSS2 in supporting histone acetylation and global transcriptional regulation cannot be ignored. It is likely that manipulating ACSS2 activity causes shifts in epigenetic patterns, which could have a much more global impact on biological processes and signaling pathways. Future studies will be aimed at dissecting the role of ACSS2 in epigenetic maintenance in ccRCC.

Exploration into the clinically translatable effects of pharmacologically targeting ACSS2 has yet to commence. The results of our study demonstrate ACSS2 inhibition is a well-tolerated and effective strategy to ablate HIF-2α expression and signaling in pre-clinical models of ccRCC. Furthermore, when we employed a cell line known to be resistant to HIF-2α inhibition, targeting ACSS2 proved capable of overcoming this resistance. This is the first study in ccRCC to highlight ACSS2 as an integral mediator of HIF-2α protein stability likely by regulating the post-translational modification landscape and MUL1-directed degradation. These novel methods of regulating HIF-2α could provide a much-needed approach to complement and enhance the efficacy of treatment in populations resistant to HIF-2α inhibition.

## METHODS

### Materials Availability

Pharmacological agents used in this study were acquired commercially and available at Selleckchem. Mice used in this study were acquired commercially and available at Jackson Laboratories. Stable cell lines carrying targeted shRNA or with different constructs are available through establishment of Material Transfer Agreement between Vanderbilt University Medical Center and requesting institution.

### Experimental Models

#### Cell Cultures

HKC cells were a gift from L.C. Racusen. HEK293T, A498, 786-O, and 769-P cell lines were obtained from the ATCC. Prior to experimentation, all cell lines were tested for Mycoplasma contamination using the ATCC Universal Mycoplasma detection kit (ATCC, catalog number 30-1012K). HEK293T, HKC, and A498 cell lines were cultured using Gibco Dulbecco’s Modified Eagle Medium (DMEM) supplemented with 10% fetal bovine serum (FBS), 1% L-glutamine, and 10 mL/L penicillin/streptomycin (Sigma). 768-O and 769-P cell lines were cultured using Gibco RPMI-1640 supplemented with 10% FBS, 1% L-glutamine (Sigma), and 10 mL/L penicillin/streptomycin (Sigma). All cells were maintained at 37°C in a 5% CO2 incubator.

#### Animal Studies

6-week old male NOD-scid IL2Rg^null^ (NSG) mice were purchased from The Jackson Laboratory (catalog number 005557) and utilized in all tumorigenesis xenograft models. Mice were housed five to a cage in a temperature- and humidity-controlled space (20-25°C, 45%– 64% humidity) with regulated water and lighting (12 h light/dark) within the animal facility. All in vivo mouse studies designed and performed in accordance to animal protocols that were approved by the Institutional Animal Care and Use Committee of VUMC. Prior to inoculation, 786-O cells were cultured as described above then trypsinized, counted, and resuspended in growth media and Matrigel at a 1:1 ratio. For all in vivo experiments, NSG mice were subcutaneously injected with 100 µl of the cell suspension providing 5×10^6^ cells. Mice in pharmacological studies were then subjected to daily treatment with vehicle control or 15 mg/kg ACSS2i delivered via intraperitoneal injection. Alternatively, mice which were inoculated with 786-O cells transduced to express the pTRIPZ doxycycline-inducible shRNA system were provided a 200 mg/kg doxycycline rodent diet (Bio-Serv) and allowed to feed ad libitum. Tumor burden was monitored via weekly manual, digital caliper measurement with intermittent measurements taken as needed. Mice were euthanized once the first tumor reached size endpoint (1,000 mm^3^) or if discomfort was observed as outlined in the IACUC approved protocol.

#### Patient Samples

Primary cell cultures were derived from tumors of patients diagnosed with clear cell renal cell carcinoma being treated at Vanderbilt University Medical Center. Informed written consent was collected from all patients whose samples were used in this study and all samples were processed and utilized in accordance with the IRB protocol (151549). Single cell suspensions were generated from the tumors and were subsequently cultured in Gibco RPMI-1640 media supplemented with 10% FBS, 1% L-Glutamine (Sigma), B-27 supplement (Gibco; 10 mL, 50x stock), and 1x Antibiotic-Antimycotic (Gibco). Fresh growth media was provided every other day for approximately 10 days until a monolayer of epithelial cells was established, at which point the primary cells were able to be passaged.

### Method Details

#### Protein Extraction and Western Blotting

Cells were harvested and suspended in RIPA buffer supplemented with 1x Halt protease inhibitor cocktail (ThermoFisher) and subjected to mechanical needle lysing with a 25G needle (BD) while being kept on ice. Cell lysates were pelleted via centrifugation at 14,000 x g for 5 minutes at 4°C and supernatants were transferred to fresh Eppendorf tubes for immediate analysis via SDS-PAGE or preservation at −20°C. Protein concentrations were determined using the BCA protein quantification method. For SDS-PAGE analysis, 50 µg of protein per sample was loaded into and run on a 4-20% gradient polyacrylamide gel (Bio-rad), followed by transfer onto a PVDF membrane. The membranes were then subjected to immunoblotting with the primary antibodies found in **Supplemental Table 1**. For western blot analysis, β-Actin served as the loading control.

#### Immunoprecipitation

Cell lysates were prepared as described above and 500 µg of total protein per sample was first cleared via incubation with protein A agarose beads at 4°C for 1 hour then spun down at 3,500 x g for 10 minutes at 4°C. The pre-cleared lysates were then transferred to fresh Eppendorf tubes and immunoprecipitation commenced via incubation with the primary antibody overnight at 4°C. The following day, 30 µl of a 50% protein A agarose bead slurry was added to the immunoprecipitation reactions and continued incubating for 2 hours at 4°C on a rotator. The immunoprecipitations were then microcentrifuged at 3,500 x g for 10 minutes at 4°C to collect the beads, which were then washed five times with Ripa buffer supplemented with protease inhibitors. After the final wash, the pelleted beads were resuspended with 3X SDS sample buffer, vortexed briefly, then microcentrifuged at 14,000 x g for 1 minute at room temperature. Samples were then heated at 95°C for 5 minutes, microcentrifuged at 14,000 x g for 1 minute at room temperature and loaded into a gel for SDS-PAGE to then be analyzed via western blotting.

#### Lentiviral Transduction

HEK293T cells were used for packaging the vectors in Supplemental Table 2 into lentivirus. HEK293T cells were allowed to reach 70% confluency and on the day of transfection, 1 hour prior to transfection growth media was replaced with Optimem. For each plate, a transfection cocktail was was prepared using 20 µg of the experimental plasmid (**Supplemental Table 2**), 15 µg PAX2, 6.5 µg of pMD2.G, 62 µl of CaCl_2_, and sterile dH_2_O to bring the total volume to 500 µl, then 500 µl of 2X HBS was added and mixed by pipetting. The transfection media was then added to the HEK293T cells, gently swirled, and incubated for 5-7 hours at 37°C. Transfection medium was then discarded and replaced with 10-12 mL of fresh growth media and the cells were allowed to produce viral particles for 48-72 hours. The lentiviral supernatants were then collected and passed through a 0.45 micron filter to be used immediately or stored at −80°C for later use.

Lentiviral transduction of 786-O cells was performed by mixing 1 mL lentivirus, 6.5 µl of 10 mg/mL polybrene, and 7 mL growth media and incubating for 24 hours at 37°C. The next day, transduction media was replaced with fresh growth media and cells were returned to 37°C incubator for another 24 hours. On the following day, the media of the transduced cells was supplemented with the antibiotic corresponding to the selectable antibiotic resistance marker in each plasmid for 24 hours, and the resistant clones were expanded for experimental use.

#### RNA Isolation and qRT-PCR

Total RNA extracts were isolated and purified using the RNeasy RNA isolation kit (Qiagen) and converted to cDNA via reverse transcription (Applied Biosystems, High Capacity cDNA Reverse Transcription kit). For each experiment, 25 ng of cDNA per sample was combined with Taqman Universal Master Mix (Applied Biosystems) and RNAse-free water in a 96-well reaction plate and qRT-PCR was performed using a Bio-Rad CFX96 thermocycler. Primers used for qRT-PCR reactions were Epas1 (Hs01026149_m1), HKII (Hs00606086_m1), SLC2A1 (Hs00892681_m1), and 18S rRNA (Hs99999901_s1). Data analysis was performed using CFX Manager software (Bio-Rad) and transcript abundance was determined via the ΔΔCt with normalization to 18S rRNA.

#### Metabolic Gene Expression

To analyze changes in expression of genes involved in metabolism, we extracted RNA as described above and performed a gene expression assay using the Nanostring nCounter Metabolic Pathways panel. The assay was performed by Vanderbilt Advanced Technologies for Genomics (VANTAGE) core facility following the manufacturer’s recommended procedures. RNA concentrations were determined via Qubit and normalized to 20ng/ul. For each sample hybridization reaction, we used 100 ng of total RNA and hybridizations were performed for a total of 20 hours. Analysis of raw data was performed using the nSolver Analysis Software 4.0 (Nanostring). For all analyses, the background threshold parameters were set to the mean of 8 negative control spike-in genes and mRNA expression was normalized to the geometric mean of 6 positive control spike-in genes and 20 housekeeping genes (ABCF1, AGK, COG7, DHX16, DNAJC14, EDC3, FCF1, G6PD, MRPS5, NRDE2, OAZ1, POLR2A, SAP130, SDHA, STK11IP, TBC1D10B, TBP, TLK2, UBB, and USP39).

Data was analyzed by ROSALIND® (https://rosalind.bio/), with a HyperScale architecture developed by ROSALIND, Inc. (San Diego, CA). Read Distribution percentages, violin plots, identity heatmaps, and sample MDS plots were generated as part of the QC step. The limma R library was used to calculate fold changes and p-values and perform optional covariate correction. Clustering of genes for the final heatmap of differentially expressed genes was done using the PAM (Partitioning Around Medoids) method using the fpc R library that takes into consideration the direction and type of all signals on a pathway, the position, role and type of every gene, etc. Hypergeometric distribution was used to analyze the enrichment of pathways, gene ontology, domain structure, and other ontologies. The topGO R library, was used to determine local similarities and dependencies between GO terms in order to perform Elim pruning correction. Several database sources were referenced for enrichment analysis, including Interpro, NCBI, MSigDB, REACTOME, WikiPathways. Enrichment was calculated relative to a set of background genes relevant for the experiment.

#### Crystal Violet Cell Growth Assay

To assess the impact of the experimental conditions on cell growth, 2.5 x 10^4^ or 1 x 10^5^ cells were seeded into 12-well or 6-well plates, respectively, and allowed to adhere for 24 hours. On the following day, growth media was supplemented with vehicle, 1 µg/mL doxycycline, or various doses of the ACSS2 inhibitor and the effect on growth was monitored over time by staining with a 1% crystal violet solution prepared in 20% methanol and 80% dH_2_O. After imaging, the crystal violet stain was then stripped using a 1% deoxycholate solution, which was transferred to a 96-well plate and read on a Promega GloMax plate-reader at 560 nm to quantify the crystal violet staining.

#### BrdU Cell Proliferation Assay

To determine the impact of experimental conditions on proliferation, we used a BrdU cell proliferation ELISA kit (Abcam, catalog number ab126556) following the manufacturer’s suggested protocol. Briefly, cells were seeded at a density of 1 x 10^4^ cells/well in normal growth media. Additionally, a duplicate set of cells to those in the experiment were seeded and did not receive BrdU labeling reagent, while several wells were also left absent of cells to thoroughly account for background. For 24-hour measurements, growth media was supplemented with ACSS2 inhibitor or doxycycline, where applicable, and allowed to incubate for 22 hours at 37°C. BrdU labeling solution was then added for the final 2 hours of the 24-hour incubation before continuing the ELISA protocol. For 48-hour measurements growth media was supplemented with ACSS2 inhibitor and incubated for 24 hours at 37°C before adding the BrdU labeling solution for the final 24 hours of the 48-hour incubation. Following BrdU labeling, cells were fixed for 30 minutes with 1X fixing solution, washed 3 times with 1X wash buffer, and then incubated with detector antibody for 1 hour at room temperature. After incubation with the detector antibody, cells were washed 3 times with 1X wash buffer followed by the addition of 1X Peroxidase Goat Anti-Mouse IgG Conjugate for 30 minutes at room temperature. The ELISA reactions were washed 3 more times with 1X wash buffer and a final time with distilled water. After patting plates dry, TMB Peroxidase substrate was added and reactions incubated for 30 minutes at room temperature in the dark. The reactions were stopped via the addition of stop solution and BrdU incorporation was quantified by reading the plates at 450 nm on a GloMax plate-reader.

#### Tumor Sphere Formation Assay

786-O cells were seeded onto 12-well ultra-low attachment plates (Corning) at a density of 1 x 10^3^ cells per well. Cells were subjected to treatment immediately following plating and fresh growth media with vehicle or the ACSS2 inhibitor was provided every other day. Sphere formation was monitored via microscope daily and after 7 days, images were captured at 4X and 10X magnification on a Keyence BZ-X800 microscope.

#### Anchorage-Independent Growth Assay

Tumorigenic potential was evaluated using an anchorage-independent growth assay. Briefly, a 2.5% agarose solution was prepared using low-melt agarose and autoclaved dH_2_O. This solution was diluted 1:4 in growth media and used to coat 6-well plates which were returned to the 37°C incubator and allowed to set, while cells were trypsinized and counted. For each condition, 1 x 10^5^ cells were resuspended in 9 mL of growth media and 1 mL of a 3% agarose solution (0.3% final concentration) and 1 mL (1 x 10^4^ cells) of the cell solution was added per well. The top layer was allowed to set for an hour at 37°C at which point growth 2 mL growth media was added to each well. Growth medium and treatments, where applicable, were exchanged every other day and colony formation was monitored via microscope. Images of colonies were captured using EVOS imaging system.

#### Immunohistochemistry Staining and Imaging

Tumor specimens were harvested immediately following animal euthanasia. Each tumor was washed with cold PBS and then fixed overnight with 10% neutral buffered formalin. The tumors were then paraffin-embedded, sectioned, and stained by the Translational Pathology Shared Resource facility at VUMC using standard IHC procedures. Images of stained tumor sections were obtained using an Olympus light microscope and “name” imaging software.

#### HIF-2α Protein Stability Assay

To assess the impact of ACSS2 inhibition on HIF-2α protein stability, 786-O cells were treated with 1 µM of the proteasomal inhibitor MG132 for 1 hour. After the initial incubation with MG132, growth media was exchanged and supplemented with DMSO or the ACSS2 inhibitor for 24 hours. Upon completion of the treatments, cell lysates were prepared as described above and analyzed via SDS-PAGE and western blotting. HIF-2α expression was quantified using ImageJ and normalized to β-Actin.

### GC/MS for Cholesterol Quantification

Lipids were extracted using a previously described method [54]. Briefly, the extracts were filtered, and lipids recovered in the chloroform phase. Individual lipid classes were separated by thin layer chromatography using Silica Gel 60 A plates developed in petroleum ether, ethyl ether, acetic acid (80:20:1) and visualized by rhodamine 6G. Phospholipids, diglycerides, triglycerides and cholesteryl esters were scraped from the plates and methylated using BF3 /methanol as described previously [55]. The methylated fatty acids were extracted and analyzed by gas chromatography. Gas chromatographic analyses were carried out on an Agilent 7890A gas chromatograph equipped with flame ionization detectors, a capillary column (SP2380, 0.25 mm x 30 m, 0.25 µm film, Supelco, Bellefonte, PA). Helium was used as a carrier gas. The oven temperature was programmed from 160 °C to 230 °C at 4 °C/min. Fatty acid methyl esters were identified by comparing the retention times to those of known standards. Inclusion of lipid standards with odd chain fatty acids permitted quantitation of the amount of lipid in the sample. Dipentadecanoyl phosphatidylcholine (C15:0), diheptadecanoin (C17:0), trieicosenoin (C20:1), and cholesteryl eicosenoate (C20:1) were used as standards. For total cholesterol, internal standard (5-a-cholestane) was added to a portion of the lipid extract and then saponified at 80 °C in 1 N KOH in 90% methanol for 1 hour. The nonsaponifiable sterol was extracted into hexane, concentrated under nitrogen, and then solubilized in hexane to inject onto the gas chromatograph. For unesterified cholesterol, internal standard is added to a portion of the lipid extract, concentrated under nitrogen and then solubilized in hexane to inject onto the gas chromatograph. The Agilent 7890A gas chromatograph was equipped with an HP-50+ column (0.25 mm i.d x 30 m, Agilent) and a flame ionization detector. The oven temperature was programmed from 260 °C to 280 °C and helium was used as the carrier gas [56].

#### Acetate Quantitation Assay

To monitor ACSS2 activity, acetate concentrations in the growth media were measured over time using an acetate quantitation colorimetric assay (BioVision, catalog number K658) using the manufacturer’s suggested protocol. Briefly, cells were seeded at a density of 2.5 x 10^5^ cells/well in a 6-well plate. The following day fresh growth media was added to the cells, as well as a blank well, supplemented with ACSS2i or doxycycline, where applicable, and incubated for 24 hours at 37°C. Growth media was collected after the 24-hour incubation and immediately processed for analysis with the colorimetric assay. For each sample, 25 µl of media was combined with 25 µl of assay buffer and 50 µl of the reaction buffer using the manufacturer provided reagents and guidelines. Following a 40-minute incubation at room temperature, acetate concentration measurements were obtained via colorimetric detection at OD450 nm using a GloMax plate-reader.

#### Transmission Electron Microscopy

All electron microscopy reagents were purchased from Electron Microscopy Sciences. Cell cultures were grown to approximately 80% confluency, subjected to treatment conditions, and fixed with 2.5% glutaraldehyde in 0.1 M cacodylate for 1 hour at room temperature followed by 24 hours at 4°C. After fixation the cells were mechanically lifted from the tissue culture plates and pelleted then sequentially post-fixed with 1% tannic acid, 1% OsO4, and en bloc stained with 1% uranyl acetate. Samples were dehydrated in a graded ethanol series, infiltrated with Quetol 651 based Spurr’s resin, and polymerized at 60°C for 48 hours. Ultrathin sections were prepared on a UC7 ultramicrotome (Leica) with a nominal thickness of 70 nm and collected onto 300 mesh nickel grids. Sections were stained with 2% uranyl acetate and lead citrate.

Samples were imaged using a Tecnai T12 operating at 100 kV equipped with an AMT NanoSprint CMOS camera using AMT imaging software. Analysis of the TEM data was performed in FIJI.

#### Mitochondrial Biogenesis Assay

The synthesis of mitochondria was assessed using a Mitochondrial Biogenesis In-Cell ELISA colorimetric kit (Abcam, catalog number ab110217), which measures the abundance of subunit I of complex IV (mtDNA-encoded) and the 70 kDa subunit of complex II (nuclear-encoded). For each experiment, the manufacturer’s suggested procedures were followed. Briefly, 1.5 x 10^4^ cells were seeded into the wells of a 96-well plate and subjected to treatments 24 hours later. Upon completion of the experimental conditions, cells were fixed for 20 minutes with 4% paraformaldehyde followed by 3 washed with PBS and the plates were then blotted dry. Subsequently, 100 µl of 0.5% acetic acid was added to each well for 5 minutes to block endogenous alkaline phosphatase activity. The wells were then washed once again with PBS followed by the addition of 1X permeabilization buffer for 30 minutes. Following permeabilization, cells were incubated with 1X blocking solution for 2 hours followed by overnight incubation with primary antibody at 4°C. After incubation with primary antibody, cells were washed 3 times with PBS and then incubated with secondary antibody for 1 hour at room temperature. Secondary antibody was then removed and the cells were washed 4 times with 1X wash buffer. Expression of the 70 kDa subunit of complex II (SDH-A) and subunit I of complex IV (COX-I) was then detected and quantified by wavelength detection at 405 nm and 600 nm, respectively, using a GloMax plate reader.

#### UALCAN and GEPIA Databases

TCGA and CPTAC expression analyses were performed using the publicly accessible UALCAN cancer database and GEPIA database [57, 58]. The effect of MUL1 gene expression on patient survival in the kidney renal clear cell carcinoma dataset (KIRC) was determined by comparing patient outcome in low vs high expression populations. MUL1 gene expression was analyzed across individual cancer stages in the KIRC TCGA dataset and expression was statistically compared to expression in normal adjacent tissue. Mul1 protein expression in tumors was analyzed in the clear cell RCC dataset and statistically compared to expression in normal adjacent tissue.

## Author’s Contributions

Z.A.B. performed most of the experimental work, data analysis, and statistical analysis. Z.A.B., J.C.R., and W.K.R. participated in study conception and design, data interpretation, and drafting manuscript. W.A.B. and E.K.A. helped with experimental work. E.S.K. was responsible for obtaining electron microscopy images. M.W.W., R.A.H., and M.L. processed patient samples and prepared primary cell cultures for experimental work. K.E.B. provided access to patient samples. All co-authors assisted with reviewing and editing manuscript.

## Acknowledgments

The authors give special thanks to their relevant funding sources. This work was supported by AACR (WKR), R01 CA21797 (JCR) and institutional research funding from Bristol-Myers Squibb, Merck, Pfizer, Calithera Biosciences, Peloton, and Incyte. ZAB is supported by the Integrated Biological Systems Training in Oncology Ruth L. Kirschstein NRSA training grant (5T32CA119925-12). VUMC Lipid Core which is supported by DK059637 and DK020593. Electron microscopy was performed in part through the use of the Vanderbilt Cell Imaging Shared Resource (supported by NIH grants CA68485, DK20593, DK58404, DK59637 and EY08126). Immunohistochemical staining was performed in part through the use of the Translational Pathology Shared Resource at VUMC. Authors acknowledge use of an active BioRender license to generate model.

---

**Supplemental Figure 1.**
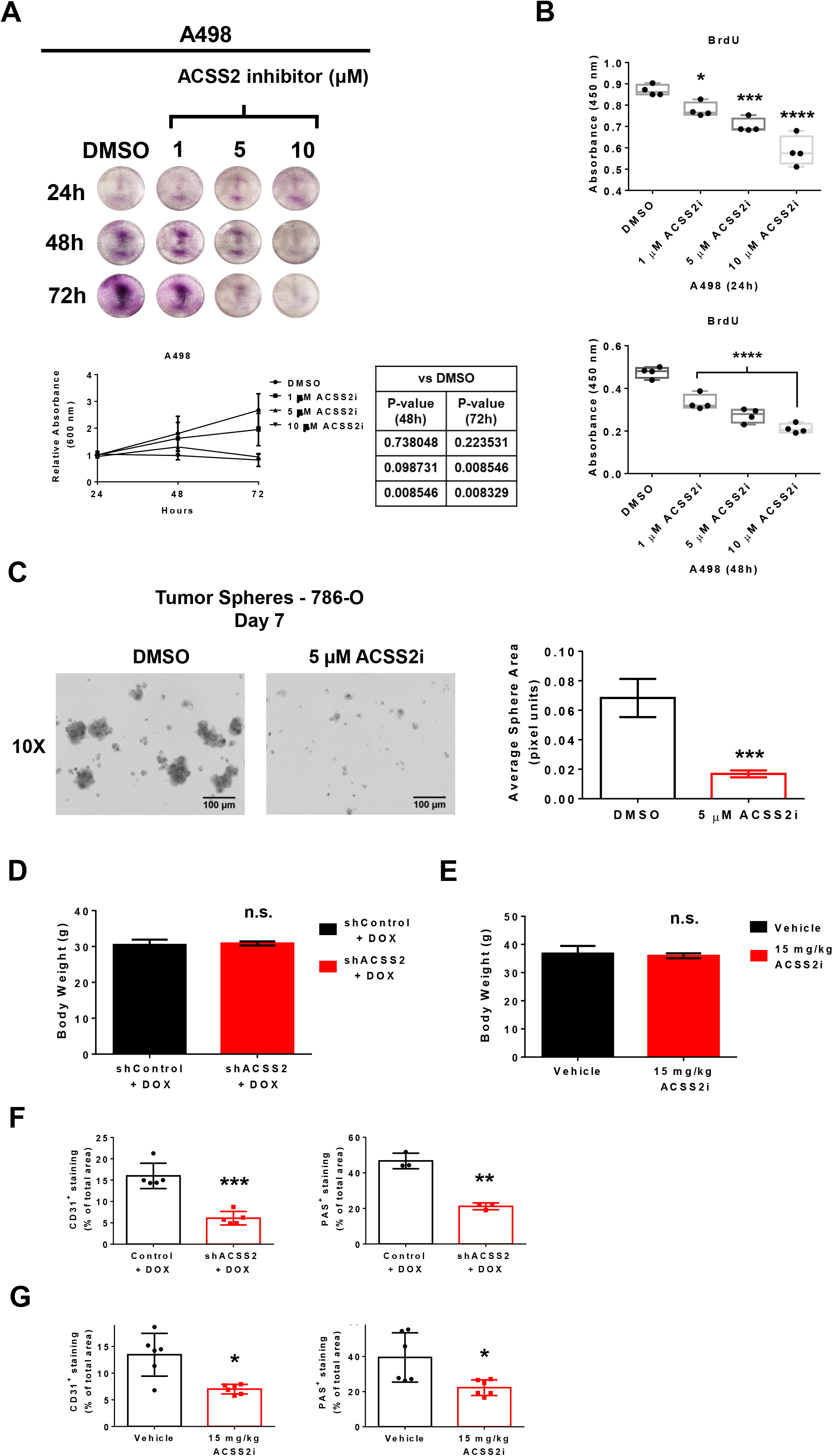
**A.** Representative images of crystal violet staining from a dose-response time-course treatment with DMSO or ACSS2 inhibitor in A498 cells. Quantification of three-independent replicates are provided in the graph below. Statistical significance was determined using Bonferroni’s multiple comparisons test. **B.** Box and whisker plots showing the absorbance values detected at OD450 nm of BrdU ELISA assays performed on A498 cells treated for 24 hours (top) or 48 hours (bottom) with DMSO, 1 µM, 5 µM, or 10 µM ACSS2 inhibitor (n=4). Statistical significance was determined using Bonferroni’s multiple comparisons test (*, P < 0.05; ***, P < 0.001; ****, P < 0.0001). **C.** Representative images of 786-O tumor spheres at day 7 of growth in ultra-low attachment plates treated with either DMSO or 5 µM ACSS2i. Quantification of average sphere area from three-independent replicates are provided in the bar graph (right). Statistical significance was determined using an unpaired, two-tailed t-test (***, P < 0.001). **D.** Bar graph showing the body weight of shControl + DOX and shACSS2 + DOX mice at the endpoint of study. **E.** Bar graph showing the body weight of Vehicle- and 15 mg/kg ACSS2i-treated mice at the endpoint of study. **F.** Bar graphs showing quantification of the percentage of total area staining positive for CD31 (left) or Periodic Acid Schiff (right) in tumor sections from shControl + DOX or shACSS2 + DOX mice. Statistical significance determined using an unpaired, two-tailed t-test with Welch’s correction (**, P-value < 0.005; ***, P-value < 0.001). **G.** Bar graphs showing quantification of the percentage of total area staining positive for CD31 (left) or Periodic Acid Schiff (right) in tumor sections from mice treated with Vehicle or 15 mg/kg ACSS2i. Statistical significance determined using an unpaired, two-tailed t-test with Welch’s correction (*, P-value < 0.05).

**Supplemental Figure 2.**
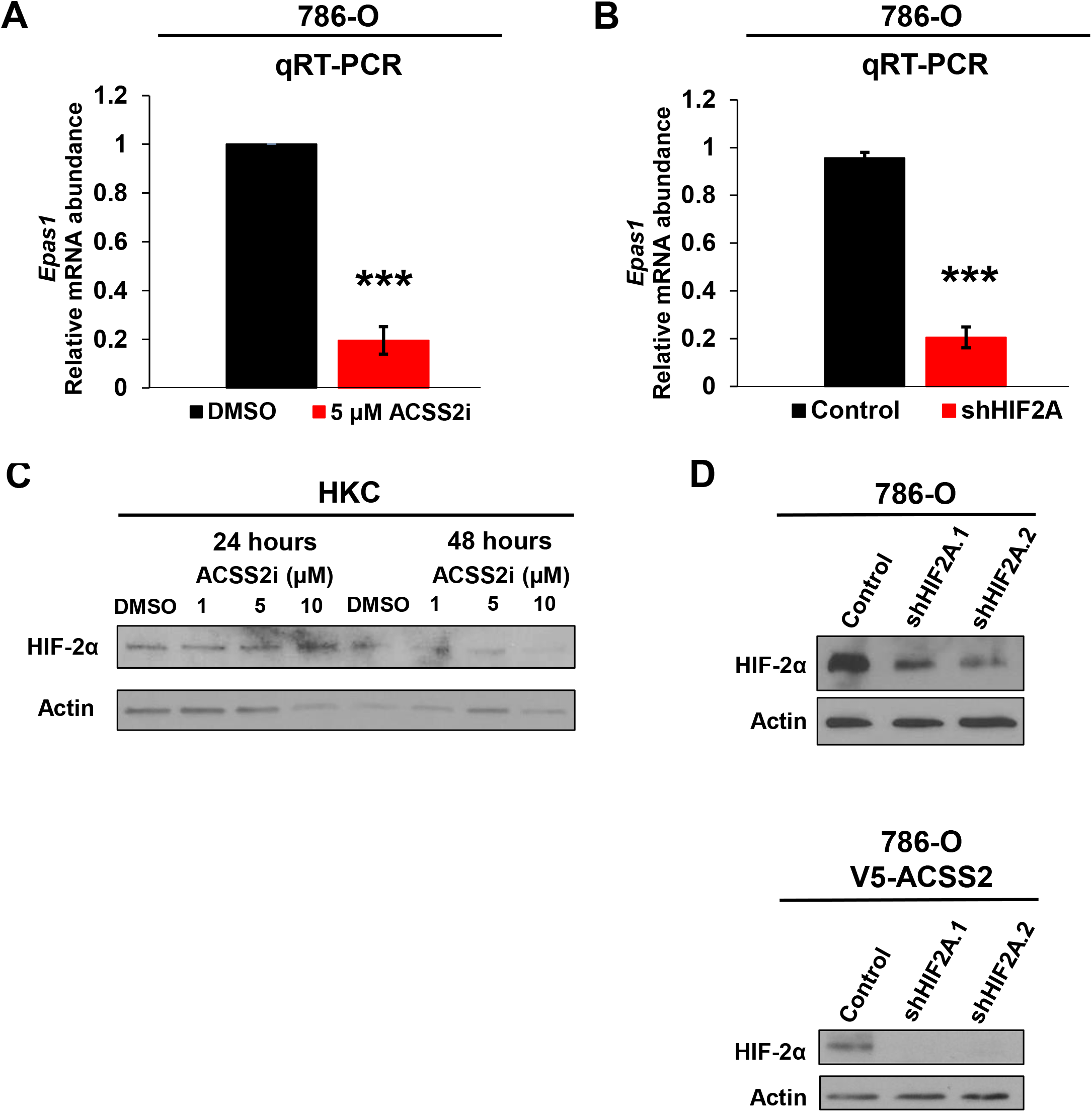
**A.** Bar graph depicting relative abundance of *Epas1* mRNA expression determined by qRT-PCR in 786-O cells treated with DMSO or 5 µM ACSS2 inhibitor for 24 hours. Statistical significance determined by unpaired, two-tailed Student’s t-test (***, P < 0.001). **B.** Bar graph depicting relative abundance of *Epas1* mRNA expression determined by qRT-PCR in 786-O cells transduced to express shControl or shHIF2A. Statistical significance determined by unpaired, two-tailed Student’s t-test (***, P < 0.001). **C.** Western blot analysis of HIF-2α and Actin in the HKC cells treated with DMSO, 1µM, 5 µM, or 10 µM ACSS2i for 24 and 48 hours.**D.** Western blot analysis of HIF-2α, ACSS2, and Actin in the 786-O cells transduced with empty vector (top) or stably overexpressing V5-ACSS2 (bottom) previously developed in Figure 2 transduced with shControl or shHIF2A targeting constructs.

**Supplemental Figure 3.**
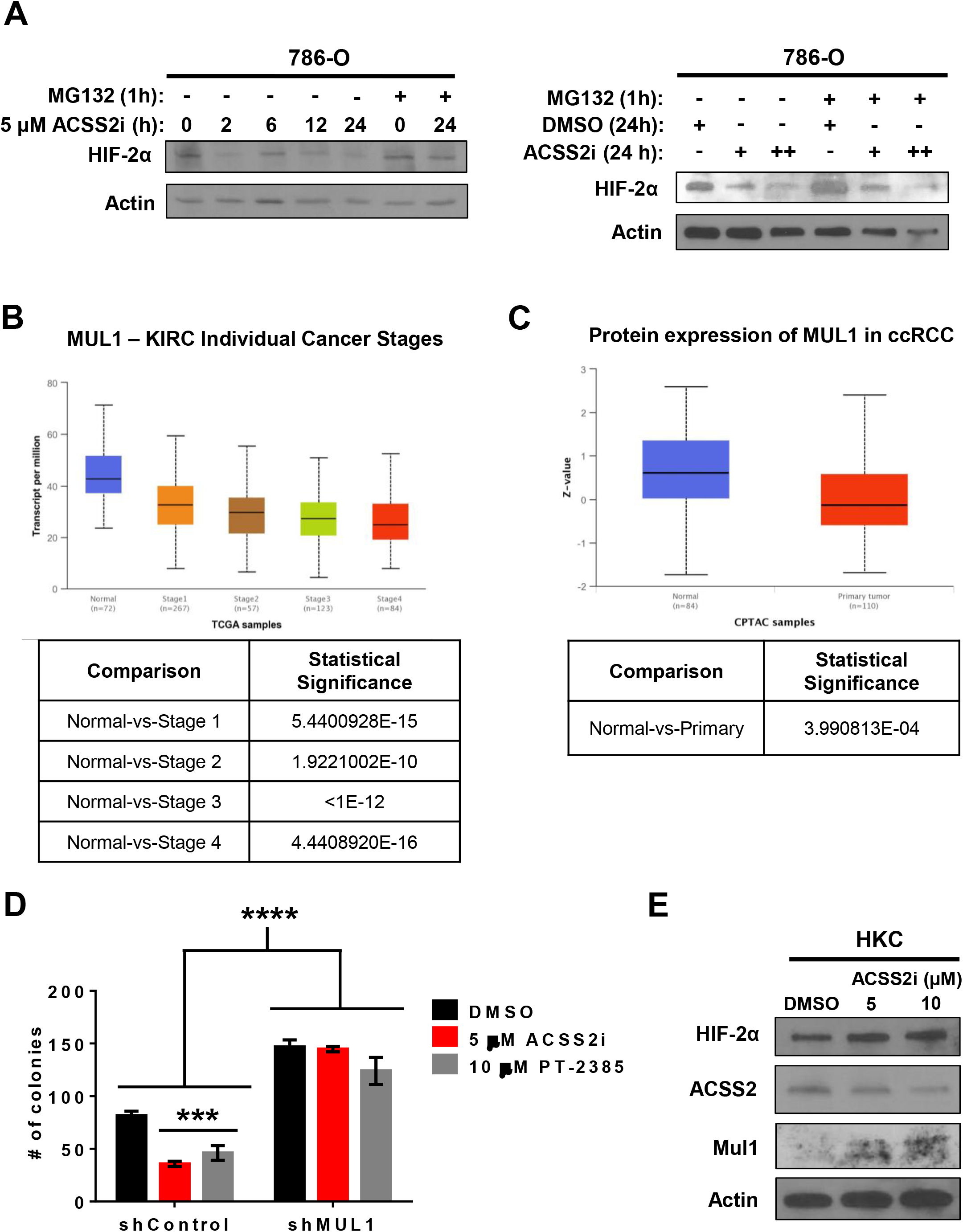
**A.** Western blot from replicate experiments depicting HIF-2α expression in 786-O cells in the absence or presence of a 1-hour incubation with 10 µM MG132 prior to being treated with DMSO or 5 µM ACSS2 inhibitor for 24 hours. **B.** Box and whisker plot generated using UALCAN database to access KIRC TCGA dataset and assess MUL1 gene expression spanning the individual stages of disease. Statistical significance determined automatically via UALCAN database. **C.** Box and whisker plot generated using UALCAN database to access KIRC CPTAC dataset and assess MUL1 protein expression in normal kidney and ccRCC samples. Statistical significance determined automatically via UALCAN database. **D.** Bar graph showing the number of colonies formed for each condition from three-independent experiments. Statistical significance determined using two-way ANOVA and Tukey’s multiple comparisons test (***, P-value < 0.0005; ****, P-value < 0001). **E.** Western blot analysis of HIF-2α, ACSS2, MUL1, and Actin in the HKC cells treated with DMSO, 5 µM, or 10 µM ACSS2i for 24 hours.

**Supplemental Figure 4.**
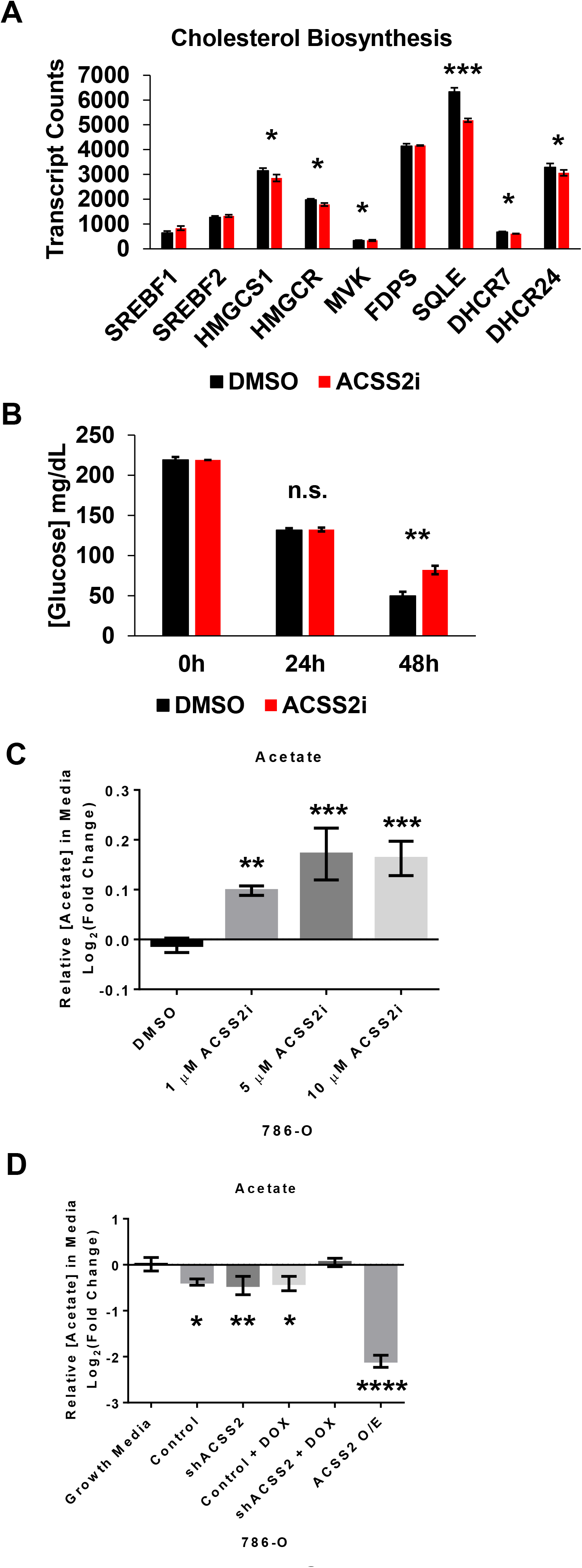
**A.** Bar graph showing transcript counts for genes involved in the cholesterol biosynthesis pathway extracted from the NanoString dataset developed from the experiment in Figure 4A where 786-O cells were treated with DMSO (black) or 5 µM ACSS2i (red) for 24 hours. Statistical significance determined by performing multiple unpaired, two-tailed t-tests (*, P-value < 0.05; ***, P-value < 0.0005). **B.** Bar graph showing average glucose concentrations in growth media in 786-O cells treated with DMSO (black) or 5 µM ACSS2i (red) at 0, 24, and 48 hour time points. Statistical significance determined using unpaired, two-tailed t-test (***, P-value < 0.005). **C.** Bar graph showing relative concentrations of acetate in growth media of 786-O cells treated with DMSO, 1 µM, 5 µM, or 10 µM ACSS2 inhibitor for 24 hours. Data are presented as Log₂ values of fold change. Statistical significance determined using an ordinary one-way ANOVA and Bonferroni’s multiple comparisons test (**, P-value < 0.01; ***, P-value < 0.001). **D.** Bar graph showing relative concentrations of acetate in growth media of 786-O cells transduced to express V5-ACSS2 overexpression vector, or the DOX-inducible shControl or shACSS2 constructs. Data are presented as Log₂ values of fold change. Statistical significance determined using an ordinary one-way ANOVA and Bonferroni’s multiple comparisons test (*, P-value < 0.05; **, P-value < 0.01; ****, P-value < 0.0001).

**Supplemental Figure 5.**
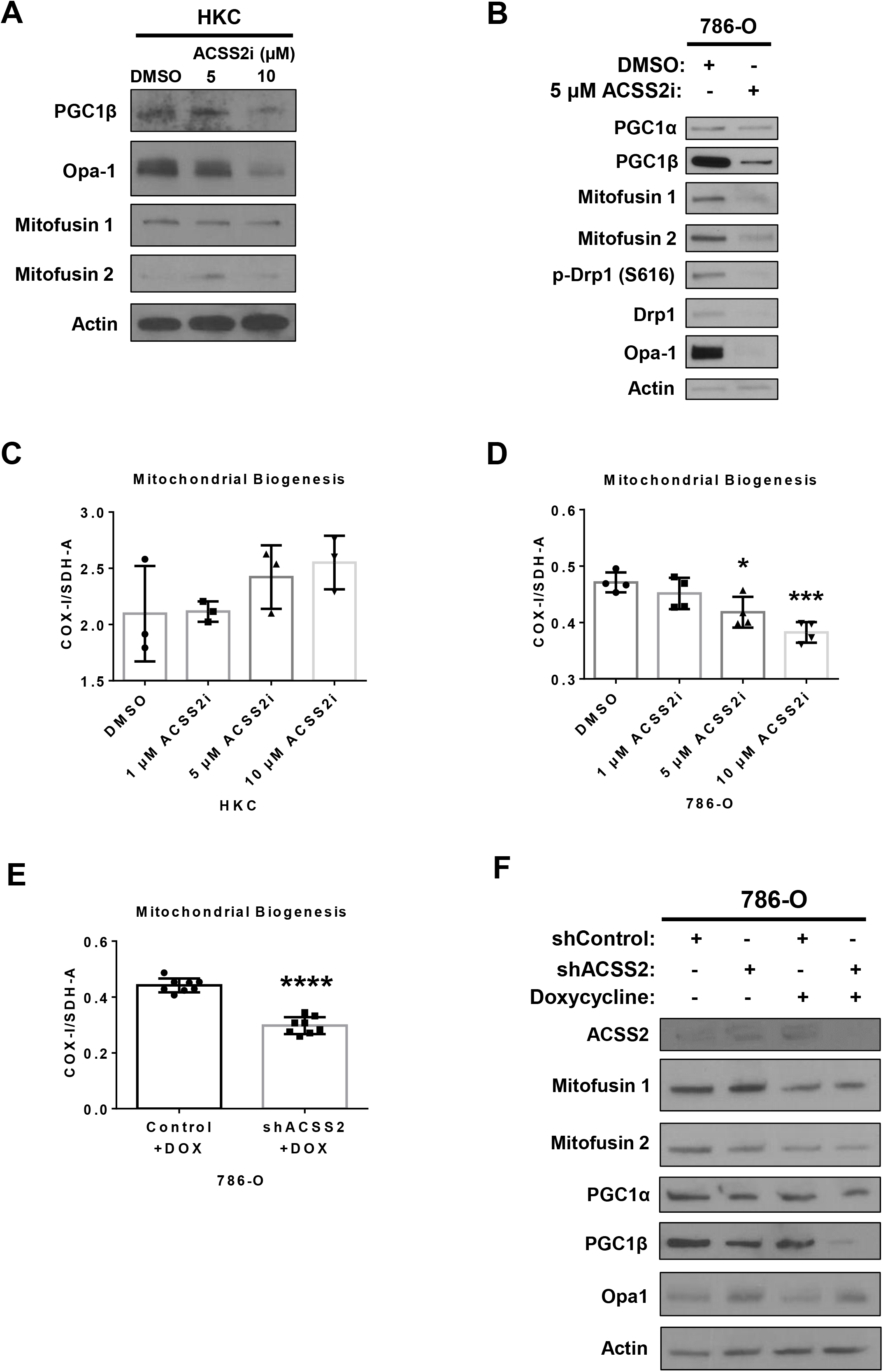
**A.** Western blot analysis of PGC1β, Opa-1, Mitofusin 1, Mitofusin 2, and Actin using the same membrane from Supplemental Figure 3E. Experiment was performed in HKC cells treated with DMSO, 5 µM or 10 µM ACSS2 inhibitor for 24 hours. **B.** Western blot analysis of PGC1α, PGC1β, Opa-1, Mitofusin 1, Mitofusin 2, Drp1, phospho-Drp1 (S616), and Actin using same membrane as Figure 3D. Experiment performed in 786-O cells treated with DMSO or 5 µM ACSS2 inhibitor for 24 hours. **C.** Bar graph with individual data points showing quantification of mitochondrial biogenesis in HKC cells treated with DMSO, 5 µM or 10 µM ACSS2 inhibitor for 24 hours. Data is shown as the ratio of expression of COX-I to SDH-A quantified by wavelength detection at 405 nm (SDH-A) and 600 nm (COX-I). Statistical significance assessed using an ordinary one-way ANOVA and Bonferroni’s multiple comparisons test. **D.** Bar graph with individual data points showing quantification of mitochondrial biogenesis in 786-O cells treated with DMSO, 5 µM or 10 µM ACSS2 inhibitor for 24 hours. Data is shown as the ratio of expression of COX-I to SDH-A quantified by wavelength detection at 405 nm (SDH-A) and 600 nm (COX-I). Statistical significance determined using an ordinary one-way ANOVA and Bonferroni’s multiple comparisons test (*, P-value < 0.05; ***, P-value < 0.001). **E.** Bar graph with individual data points showing quantification of mitochondrial biogenesis in 786-O pTRIPZ shControl and shACSS2 cells treated with doxycycline for 24 hours. Data is shown as the ratio of expression of COX-I to SDH-A quantified by wavelength detection at 405 nm (SDH-A) and 600 nm (COX-I). Statistical significance determined using an unpaired, two-tailed t-test (****, P-value < 0.0001). **F.** Western blot analysis of PGC1α, PGC1β, Opa-1, Mitofusin 1, Mitofusin 2, and Actin probing the same membrane as used in Figure 3E. Experiment performed in 786-O pTRIPZ shControl and shACSS2 cells treated with doxycycline for 24 hours.

**Supplemental Figure 6.**
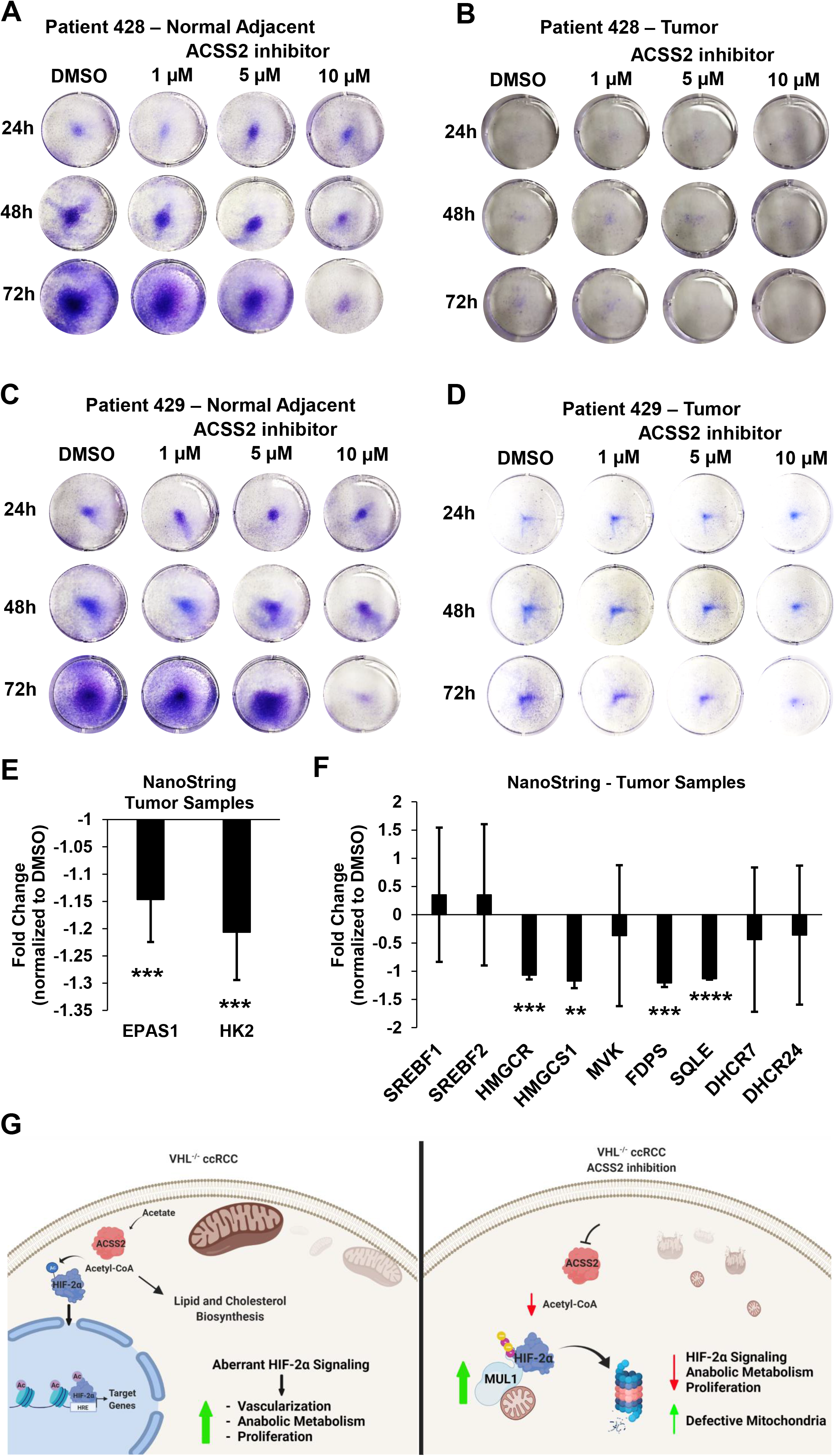
**A.** Representative images of crystal violet staining from a dose-response time-course treatment with DMSO or ACSS2 inhibitor in epithelial cells isolated from ccRCC patient’s normal adjacent tissue. **B.** Representative images of crystal violet staining from a dose-response time-course treatment with DMSO or ACSS2 inhibitor in epithelial cells isolated from ccRCC patient tumor sections. **C.** Representative images of crystal violet staining from a dose-response time-course treatment with DMSO or ACSS2 inhibitor in epithelial cells isolated from ccRCC patient’s normal adjacent tissue. **D.** Representative images of crystal violet staining from a dose-response time-course treatment with DMSO or ACSS2 inhibitor in epithelial cells isolated from ccRCC patient tumor sections. **E.** Bar graph showing average fold change values for gene expression of *HK2* and *EPAS1* extracted from Nanostring experiment performed in ccRCC patient tumor samples treated with DMSO (n=3) or 5 µM ACSS2i (n=3) for 48 hours. Statistical significance determined by performing unpaired, two-tailed t-tests (***, P-value < 0.001). **F.** Bar graph showing average fold change values for gene expression of genes involved in the cholesterol biosynthesis pathway extracted from Nanostring experiment performed in ccRCC patient tumor samples treated with DMSO (n=3) or 5 µM ACSS2i (n=3) for 48 hours. Statistical significance determined by performing multiple unpaired, two-tailed t-tests (**, P-value < 0.01; ***, P-value < 0.001; ****, P-value < 0.0001). **G.** Model of mechanism.

