## Supplemental Tables 1 and 2 for "ACSS2 Regulates HIF-2α Degradation through the E3-Ubiquitin Ligase MUL1 in Clear Cell Renal Cell Carcinoma"

**TABLE LEGENDS**

**Supplemental Table 1.** List of primary antibodies used throughout the study.

**Supplemental Table 2.** List of plasmid constructs used throughout the study.

**TABLES**

**Supplemental Table 1**

| **Antibody** | **Application** | **Concentration** | **Source** | **Catalog Number** |
| --- | --- | --- | --- | --- |
| ACSS2 | WB | 1:1,000 | Cell Signaling | 3658S |
| Β-Actin | WB | 1:5,000 | Abcam | ab8227 |
| Drp1 | WB | 1:1,000 | Cell Signaling | 8570S |
| p-Drp1 (S616) | WB | 1:1,000 | Cell Signaling | 3455S |
| EGLN3 | WB | 1:1,000 | Novus | NB100-139 |
| EPAS1 | WB, IP | 1:500, 1:50 | Santa Cruz | sc-46691 |
| Epo | WB | 1:1,000 | Santa Cruz | sc-5290 |
| GLUT1 | WB | 1:1,000 | Abcam | ab115730 |
| HIF-2α | WB, IP | 1:500, 1:100 | Novus | NB100-122 |
| Hexokinase II | WB | 1:1,000 | Cell Signaling | 2867S |
| K48 polyubiquitin | WB | 1:1,000 | Cell Signaling | 8081S |
| LC3 A/B | WB | 1:1,000 | Cell Signaling | 12741S |
| Mitofusin 1 | WB | 1:1,000 | Cell Signaling | 14739S |
| Mitofusin 2 | WB | 1:1,000 | Cell Signaling | 11925S |
| MUL1 | WB, IP | 1:500; 1:100 | Abcam | ab84067 |
| Opa1 | WB | 1:1,000 | Cell Signaling | 80471S |
| p62 | WB | 1:1,000 | Cell Signaling | 5114S |
| PGC1α | WB | 1:1,000 | Novus | NBP1-04676 |
| PGC1β | WB | 1:1,000 | Abcam | ab176328 |
| SREBP1 | WB | 1:1,000 | Novus | NB600-582 |
| SREBP2 | WB | 1:1,000 | Novus | NBP1-54446 |
| V5-tag | WB | 1:1,000 | Cell Signaling | 13202S |
| VEGFR2 | WB | 1:500 | Cell Signaling | 9698S |

**Supplemental Table 2**

| **Plasmid** | **Clone** | **Source** | **Catalog Number** |
| --- | --- | --- | --- |
| TRIPZ Human Non-silencing shRNA | N/A | Horizon Discovery | RHS4743 |
| TRIPZ Human ACSS2 shRNA | V3THS_366720 | Horizon Discovery | RHS4696-200764456 |
| TRIPZ Human ACSS2 shRNA | V3THS_366722 | Horizon Discovery | RHS4696-200767025 |
| pLX304 CCSB-Broad LentiORF | ACSS2 | Horizon Discovery | OHS6085-213584639 |
| pLX304 empty vector | N/A | Addgene | 25890 |
| GIPZ EPAS1 shRNA | V2LHS_113750,113752, 318637, 402392, 318640 | Horizon Discovery | RHS4531-EG2034 |
| pLKO MUL1 shRNA | TRCN0000033937, TRCN0000236073, | Sigma | N/A |
| pRK5-HA-Ubiquitin-WT | N/A | Addgene | 17604 |
| pRK5-HA-Ubiquitin-K48R | N/A | Addgene | 17608 |
